# Degradomic identification of membrane type 1-matrix metalloproteinase (MT1-MMP/MMP14) as an ADAMTS9 and ADAMTS20 substrate

**DOI:** 10.1101/2022.10.17.512574

**Authors:** Sumeda Nandadasa, Daniel Martin, Gauravi Deshpande, Karyn L. Robert, M. Sharon Stack, Yoshifumi Itoh, Suneel S. Apte

## Abstract

The secreted metalloproteases ADAMTS9 and ADAMTS20 are implicated in extracellular matrix (ECM) proteolysis and primary cilium biogenesis. Here, we show that clonal gene-edited RPE-1 cells in which *ADAMTS9* was inactivated, and which constitutively lack *ADAMTS20* expression, have morphologic characteristics distinct from parental RPE-1 cells. To investigate underlying proteolytic mechanisms, a quantitative N-terminomics method, terminal amine isotopic labeling of substrates (TAILS) was used to compare parental and gene-edited cells and their medium to identify ADAMTS9 substrates. Among differentially abundant N-terminally labeled internal peptides arising from secreted and transmembrane proteins, a peptide with lower abundance in the medium of gene-edited cells suggested cleavage at the Tyr^314^-Gly^315^ bond in the ectodomain of the transmembrane metalloprotease MT1-MMP, whose mRNA was also reduced in gene-edited cells. This cleavage, occurring in the MT1-MMP hinge i.e., between the catalytic and hemopexin domains, was orthogonally validated both by lack of an MT1-MMP catalytic domain fragment in the medium of gene-edited cells and restoration of its release from the cell surface by re-expression of ADAMTS9 and ADAMTS20, and was dependent on hinge *O*-glycosylation. Since MT1-MMP is a type I transmembrane protein, identification of an N-terminally labeled peptide in the medium suggested additional downstream cleavage sites in its ectodomain. Indeed, a C-terminally semi-tryptic MT1-MMP peptide with greater abundance in wild-type RPE-1 medium identified by a targeted search indicated a cleavage site in the hemopexin domain. Consistent with retention of MT1-MMP catalytic domain on the surface of gene-edited cells, pro-MMP2 activation, which requires cell-surface MT1-MMP, was increased. MT1-MMP knockdown in gene-edited ADAMTS9/20-deficient cells restored focal adhesions but not ciliogenesis. The findings expand the web of interacting proteases at the cell-surface, suggest a role for ADAMTS9 and ADAMTS20 in regulating cell-surface activity of MT1-MMP and indicate that MT1-MMP shedding does not underlie their observed requirement in ciliogenesis.

**Highlights:**

- ADAMTS9-deficient RPE-1 cells have impaired substrate attachment
- ADAMTS9 and ADAMTS20 release the MT1-MMP catalytic domain from the cell-surface
- Increased cell-surface MT1-MMP increases pro-MMP2 activation and collagenolysis
- MT1-MMP knockdown restores substrate attachment of ADAMTS9-deficient RPE-1 cells.

**In Brief:** ADAMTS9 and ADAMTS20 are homologous secreted proteases implicated in ECM proteolysis and ciliogenesis, but few relevant substrates of these proteases are currently known. Quantitative N-terminomics comparison of RPE-1 cells with *ADAMTS9* inactivation and parental RPE-1 cells identified transmembrane protease MT1-MMP (MMP14) as a novel ADAMTS9 substrate. The resulting enhanced cell-surface MT1-MMP activity in the gene-edited cells contributes to their adhesion defect, but not lack of cilia. A key physiological function of ADAMTS9/20 may be to dampen cell-surface MT1-MMP activity.

## Introduction

Secreted and cell surface proteases undertake proteolytic modification of pericellular extracellular matrix (ECM), which is relevant to cell adhesion and migration, as well as shedding of cell-surface and transmembrane proteins, whose effects include activation or inactivation of adhesion molecules, membrane-bound ligands, receptors and proteases (1). Two classes of related zinc metalloproteases (metzincins) are prominent in cell-surface proteolysis, i.e., a disintegrin-like and metalloproteinase domain (ADAM) proteases and matrix metalloproteinases (MMPs). All ADAMs are single-pass type I membrane-spanning proteins, as are four membrane-type MMPs (MT-MMPs), whereas two other cell-surface MMPs (MT4-MMP and MT6-MMP) are inserted into the cell membrane via glycosylphosphatidylinositol (GPI) anchors. Membrane localization of these proteases is likely to be a major determinant of their activity at the cell-surface and against pericellular matrix

A disintegrin-like and metalloproteinase domain with thrombo<u>s</u>pondin type 1 repeats (ADAMTS) proteases are related to both families but comprise a distinct class of secreted proteases (2). The majority of known ADAMTS substrates are ECM components (2). Some ADAMTS proteases, akin to secreted MMPs such as MMP7, have an affinity for the cell surface and/or pericellular matrix via mechanisms such as interactions with glycosaminoglycans (3), and may thus act in a cell-proximate manner. ADAMTS1 and ADAMTS6, which bind heparan-sulfate proteoglycans (4), were previously shown to cleave the syndecan-4 ectodomain (5, 6), but transmembrane proteins are generally not thought to be preferred ADAMTS substrates.

ADAMTS9 and its mammalian homolog ADAMTS20 are evolutionarily conserved proteases with readily recognizable orthologs in *C. elegans* and *D melanogaster* (7, 8). These large proteases (predicted molecular mass∼215,000-225,000 depending on glycosylation) are distinguished both by having more thrombospondin type 1 repeats than other ADAMTS proteases and by their unique C-terminal domain (the Gon-1 domain) which occurs nowhere else in the human genome/proteome (8). In mice, *Adamts9* is widely expressed from early embryonic stages on (8-13). In adult tissues, it is a product of capillary and venous endothelium and is expressed by several other cell types (10, 14). Analysis of *Adamts9* null embryos, embryos homozygous for an *Adamts9* hypomorphic gene-trap allele and of *Adamts9* conditional mouse mutants has identified essential roles for ADAMTS9 in gastrulation, cardiovascular, eye and umbilical cord development, and together with ADAMTS20, in neural tube and palate closure, interdigital web regression and skin pigmentation (10-13, 15-19). In worms, ADAMTS9 is implicated in gonad development (20, 21), in flies in tracheal, salivary gland and germ cell development (22) and in zebrafish, early germ cell migration and ovarian development (23, 24). Unexpectedly, ADAMTS9 and ADAMTS20 are necessary for formation of the primary cilium (12), a cellular organelle of post-mitotic cells that transmits signaling by hedgehog proteins and other morphogens. They are the only proteases known to have such a role. The requirement for ADAMTS9 in ciliogenesis is supported by recessive *ADAMTS9* mutations identified in humans with ciliopathies, specifically, nephronophthisis, which is a disorder of kidney development and growth that results in shrunken kidneys, and Joubert syndrome, a ciliopathy with diverse and variable clinical manifestations (25, 26). Some of the morphogenetic defects seen in *Adamts9* and *Adamts20* mouse mutants may reflect their cooperation (with each other as well as other ADAMTS proteases) in processing the chondroitin-sulfate proteoglycan versican, an important ECM component in the early to mid-gestation embryo (reviewed in (25, 26)). Versican accumulates in some tissues of *Adamts9/20* mutant embryos as a result of reduced proteolytic turnover, which was revealed using a cleavage-specific versican neoepitope antibody (12, 13). Recessive *Adamts20* mutations in mice (*Belted* (*Bt*) mutant) affect skin pigmentation (27, 28), and lead to partially penetrant soft-tissue syndactyly and cleft palate (11, 13, 16, 18), which also occur in dogs with an *ADAMTS20* variant (29). Mice with limb-specific conditional deletion of ADAMTS9 develop soft-tissue syndactyly (16). This defect is also seen in mice with elimination of a canonical ADAMTS cleavage site in the versican core protein, validating versican as an ADAMTS9 and ADAMTS20 substrate in digit separation (30). However, mice with cleavage-resistant versican lack other defects seen in *Adamts9* and *Adamts20* mutants, some of which may reflect defective cilium-mediated signaling, or reduced cleavage of other ECM substrates that are yet to be defined. Other confirmed ADAMTS9 substrates are aggrecan and fibronectin, both components of the ECM (30). A prior degradomic analysis of skin from a hypomorphic mouse *Adamts9* mutant identified several putative ECM proteins as ADAMTS9 substrates, but these remain unvalidated and their cleavage by ADAMTS20 was not previously tested (31).

The cooperative functions of ADAMTS9 and ADAMTS20 elicited using combined mouse mutants suggested that their activities may overlap. For example, impaired ciliogenesis in gene-edited RPE-1 cells lacking ADAMTS9 was restored by transfection of ADAMTS20 (12). Catalytic action of ADAMTS9 and ADAMTS20 is required for ciliogenesis (12) suggesting that cleavage of one or more shared substrates may mediate this specific role. Systematic identification of ADAMTS9 and ADAMTS20 substrates, in the ECM, at the cell-surface or in ciliogenesis pathways is therefore crucial to define the underlying molecular mechanisms of these important secreted proteases and delineate those functions which are cooperative versus those unique to each protease.

Here, we used proteomics to investigate newly observed differences in the cell-substratum interface of an ADAMTS9-deficient clonal RPE-1 line (designated as clone D12) which was generated by gene-editing of RPE-1 cells using CRISPR-Cas9 to specifically inactivate *ADAMTS9* (12). To explain the observed morphologic differences, we applied the N-terminomics strategy TAILS (32) to compare the degradomes of these cells with parental RPE-1 cells. RPE-1 cells and D12 cells constitutively lack ADAMTS20 expression, (12), maximizing the potential for discovery of ADAMTS9/20 substrates. Moreover, the D12 cells have an unequivocal ADAMTS9/20 dependent phenotype, i.e., impaired formation of the primary cilium. The results demonstrate a role for ADAMTS9 in regulating the activity of MT1-MMP, an important cell-surface metalloprotease involved in cell migration, tumor cell invasion and numerous other processes (33), including potentially, cilium length regulation.

## Results

### Perturbation of matrix contacts in ADAMTS9-deficient RPE-1 cells

Prior work had showed accumulation of ADAMTS9 substrates versican and fibronectin in the ECM of ADAMTS9 mutant D12 cells when compared to the parental RPE-1 cells (12, 34). Both substrates are relevant to cell adhesion, with versican being anti-adhesive (35-37) and fibronectin being pro-adhesive (38, 39). We therefore compared RPE-1 and D12 morphology and cell-substratum interactions using differential interference contrast (DIC) microscopy and interference reflection microscopy (IRM), an antibody-independent optical technique for visualizing cell-substratum contacts on a glass surface (40). In images obtained by IRM, areas of a cell closely adherent to the substratum appear dark whereas areas farther away from the substratum appear light (41). IRM and DIC microscopy were used both shortly (30 min) after plating and after cell spreading (24 h) to visualize the cells and their adhesive interactions. 30 min after seeding, wild-type RPE-1 cells assumed both rounded and slightly flattened shapes and many showed intense dark areas on IRM corresponding to the formation of initial adhesions with the glass substratum (Fig. 1A, Movie S1). At this time-point, D12 cells had also attached but showed few dark regions with IRM (Fig. 1A, Movie S2). After 24 h, the parental RPE-1 cells had robust fibrillar adhesions represented by discrete dark streaks whereas D12 cells specifically showed a reduction in distinct leading edge fibrillar adhesions, with an increase in leading edge close contacts (gray areas). D12 cells also showed filamentous protrusions at their trailing edges which were rarely seen in parental RPE-1 cells, and whose origin is presently unclear (Fig. 1B, Movie S3). Quantitation of cell footprint pixel intensity showed statistically significant reduction of dark pixels in D12 cells, suggestive of reduced focal/fibrillar adhesions. The differences in the cell-matrix interface at both 30 min and 24 h, and lack of primary cilia in D12 cells as previously reported (12), prompted active consideration of cell-surface and transmembrane molecules as prospective ADAMTS9 substrates. This was addressed by degradomic comparison of parental RPE-1 and D12 cells and their conditioned medium.

**Figure 1.**
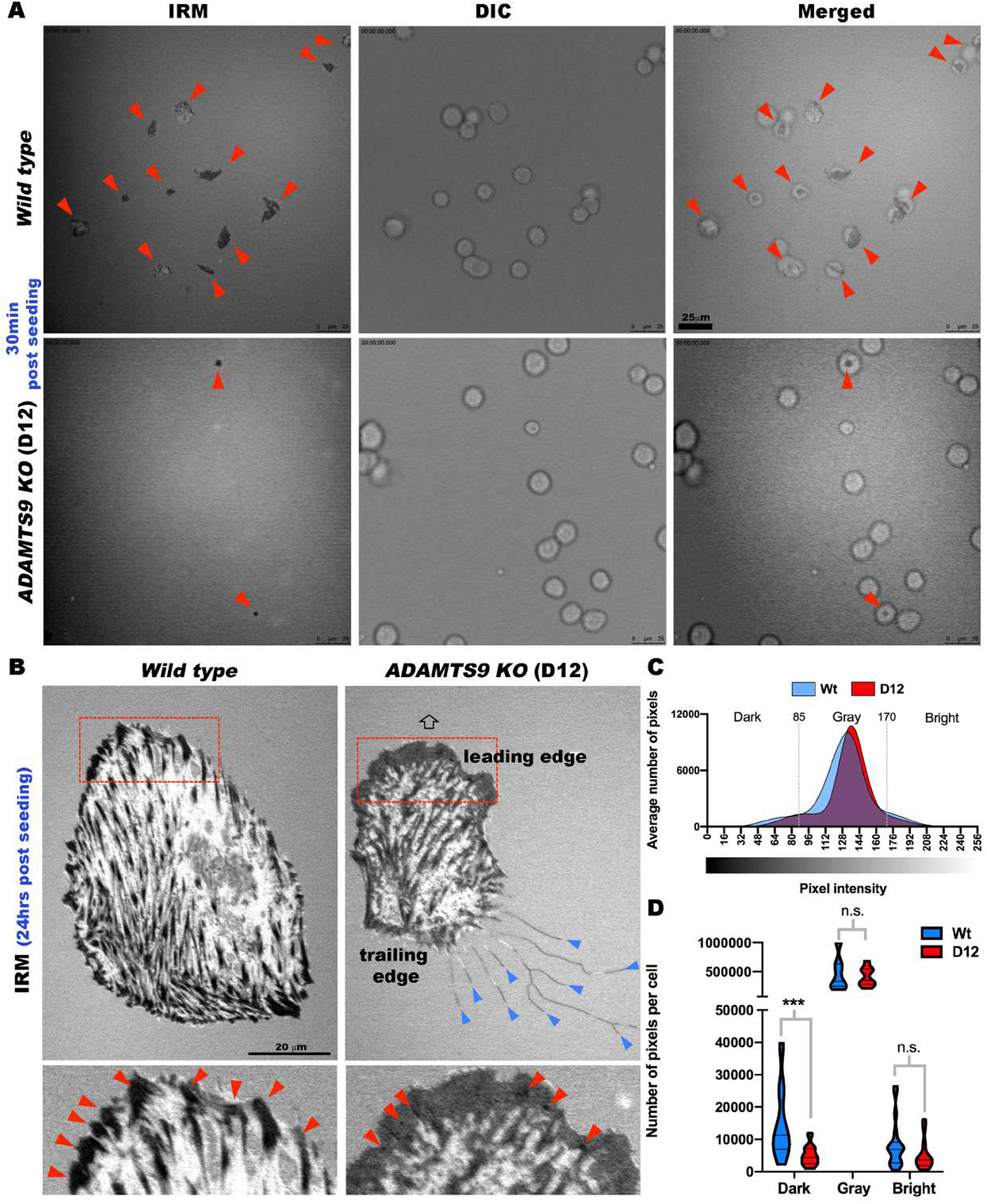
Altered cell-substratum interface in gene-edited RPE-1 cells lacking ADAMTS9. **(A)** Still photomicroscopy of concurrent, aligned interference reflection microscopy (IRM) and differential interference contrast (DIC) images from time-lapse experiments (see Supplemental Movies 1-4) at 30 minutes post-seeding show strong adhesions (dark areas in cells indicated by red arrowheads) in the majority of parental RPE-1 cells (labeled as wild type). *ADAMTS9* knockout (KO) RPE-1 cells (labeled as D12) also attach, but only occasional cells form strong adhesions at this early point. **(B)** IRM of parental RPE-1 (labeled as wild type) and *ADAMTS9* KO (D12) cells 24 hours post-seeding from time-lapse images showing poorly formed fibrillar adhesions in D12 cells (contrasting with discrete and peripheral fibrillar adhesions in parental RPE-1 cells (red arrowheads)). D12 cells show trailing edge filamentous extensions (blue arrowheads). which are lacking in parental RPE-1 cells. The arrow indicates the direction of cell migration, which was determined by live imaging. **(C)** Distribution of IRM pixel intensities comparing parental RPE-1 cells (Wt, blue) and D12 cells (red). Dotted lines bin dark (0-85), gray (85-170) and bright (170-256) pixels. **(D)** Violin plot of binned pixel intensities per cell shows that D12 cells have fewer dark pixels (strong focal adhesions). N=20 cells per group, *** p<0.0005, two-tailed unpaired Student t-test. Scale bars are 25μm in A and 20μm in B.

### TAILS identifies degradome alterations in the medium of ADAMTS9-deficient cells

Parental RPE-1 and D12 cells were cultured in serum-free medium which was analyzed using a TAILS workflow performed in triplicate, in which we employed duplex reductive dimethylation with stable light and heavy isotopes of formaldehyde for labeling the respective samples (Fig. 2A). LC-MS/MS of the differentially labeled samples, which were combined for the analysis to eliminate run-to-run variation and for precise relative quantitation, identified numerous high-confidence dimethyl-labeled N-terminal peptides. We first positionally annotated them to determine which originated internal to constitutive processing sites such as signal peptide or propeptide excision. Subsequently, statistical analysis of their relative abundance based on aligned, normalized extracted ion chromatograms (EICs) of the light/heavy MS1 peaks identified several internal peptides arising from secreted and transmembrane proteins with significantly different abundance (Fig. 2B). Among internally originating peptides (“internal peptides”) with significantly reduced abundance in the medium of D12 cells, and thus potentially indicative of ADAMTS9 substrates, were six peptides from the previously identified substrate fibronectin, a peptide from the cilium transition zone transmembrane protein Tmem67 and an N-terminally labeled MT1-MMP (MMP14) peptide, G^315^PNICDGNFDTVAMLR (Fig. 2B-D). The position of this peptide and its N-terminal dimethyl label suggested MT1-MMP cleavage at the Tyr^314^-Gly^315^ peptide bond, which is located in the so-called “hinge” between the catalytic domain and the hemopexin domain (Fig. 2E).

**Figure 2.**
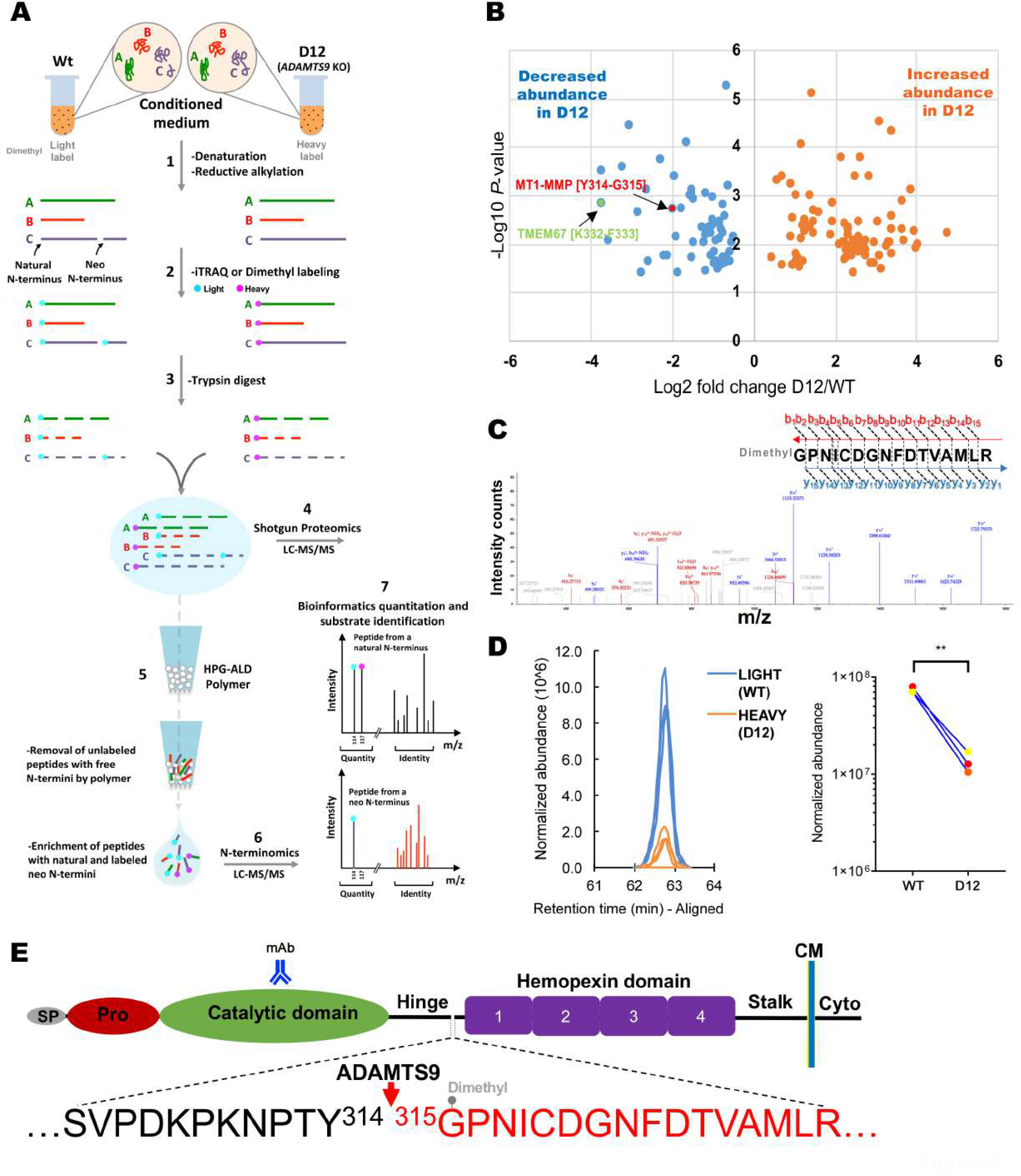
Identification of MT1-MMP as a novel ADAMTS9 substrate. **(A)** Schematic illustrating the key experimental steps of the TAILS (terminal amine isotopic labeling of substrates) strategy as applied to parental RPE-1 cells (Wt) and D12 conditioned medium (using reductive dimethylation of protein N-termini) and to cell lysates (using iTRAQ labeling of protein N-termini). **(B)** Volcano plot of positionally internal peptides arising from secreted and cell-surface proteins identified by TAILS of the medium. Peptides indicating MT1-MMP cleavage at Y^314^-G^315^ and TMEM67 cleavage at K^322^-F^333^ are identified by red and green colors respectively and show a statistically significant reduction in D12 cells. **(C)** MS2 profile of the MT1-MMP peptide (GPNICDGNFDTVAMLR) which was dimethyl labelled at the N-terminus and reduced in the medium of D12 cells. **(D)** Extracted ion chromatograms (*left*) and quantitation of normalized abundance of the identified MT1-MMP peptide in triplicate experiments (*right*). ** p<0.005, two-tailed unpaired Student t-test. **(E)** Cartoon of MT1-MMP domain structure indicating the location within the hinge of the ADAMTS9 cleavage site deduced from the peptide in (C) (SP, signal peptide; Pro, propeptide; CM, cell membrane; Cyto, cytoplasmic tail; mAb, monoclonal antibody to catalytic domain). The four blades of the of the hemopexin domain β-propeller structure are numbered in sequence from N-to C-terminus.

MT1-MMP is a type I transmembrane protein with a large ectodomain, a single membrane segment, and a very short cytoplasmic tail. Consistent with the possibility that ADAMTS9 shed the MT1-MMP ectodomain from the cell surface, western blotting under reducing conditions demonstrated a ∼35 kDa species reactive with a polyclonal antibody against the MT1-MMP catalytic domain in the medium of parental RPE-1 cells, but not the gene-edited D12 cells. (Fig. 3A). However, RT-qPCR also disclosed a statistically significant reduction in MT1-MMP (MMP14) mRNA in the D12 cells, (Fig. 3B), prompting further consideration of whether the TAILS peptide reflected reduced cleavage or reduced MT1-MMP abundance in D12 cells. Staining of RPE-1 and D12 cells without permeabilization was undertaken, which illustrated stronger MT1-MMP staining on the surface of D12 cells (Fig. 3C). Furthermore, when ADAMTS9 was co-expressed with MT1-MMP in HEK293T cells, western blotting under reducing conditions demonstrated an ∼15 kDa species as well as an additional ∼60 kDa species, both of which were absent when the catalytically inactive mutant, ADAMTS9-EA was co-expressed, validating MT1-MMP as a novel ADAMTS9 substrate (Fig. 3D). Co-expression of ADAMTS20 with MT1-MMP in HEK293T cells demonstrated the presence of a 15 kDa, but not 60 kDa species in the medium, whereas neither were seen when catalytically inactive ADAMTS20 was co-expressed (Fig. 3E). This suggested that MT1-MMP is also cleaved by ADAMTS20. Although the precise peptide bond cleaved by ADAMTS20 is unknown, the size of the catalytic domain fragment released into the medium suggested that like ADAMTS9, ADAMTS20 cleaves MT1-MMP in the hinge region (Fig. 3E). However, the discrepancy between the size of the observed band in panels A and D/E, suggests the possibility that when expressed at high levels, ADAMTS9 and ADAMTS20 may cleave at additional sites in the catalytic domain. Nevertheless, these data are firmly indicative of MT1-MMP cleavage by ADAMTS9 and ADAMTS20 and suggests release of the MT1-MMP catalytic domain from the cell-surface. The additional 60 kDa species seen upon ADAMTS9 over-expression (Fig. 3D), suggested the possibility of juxta-membrane cleavage in the MT1-MMP stalk (Fig. 3F).

**Figure 3.**
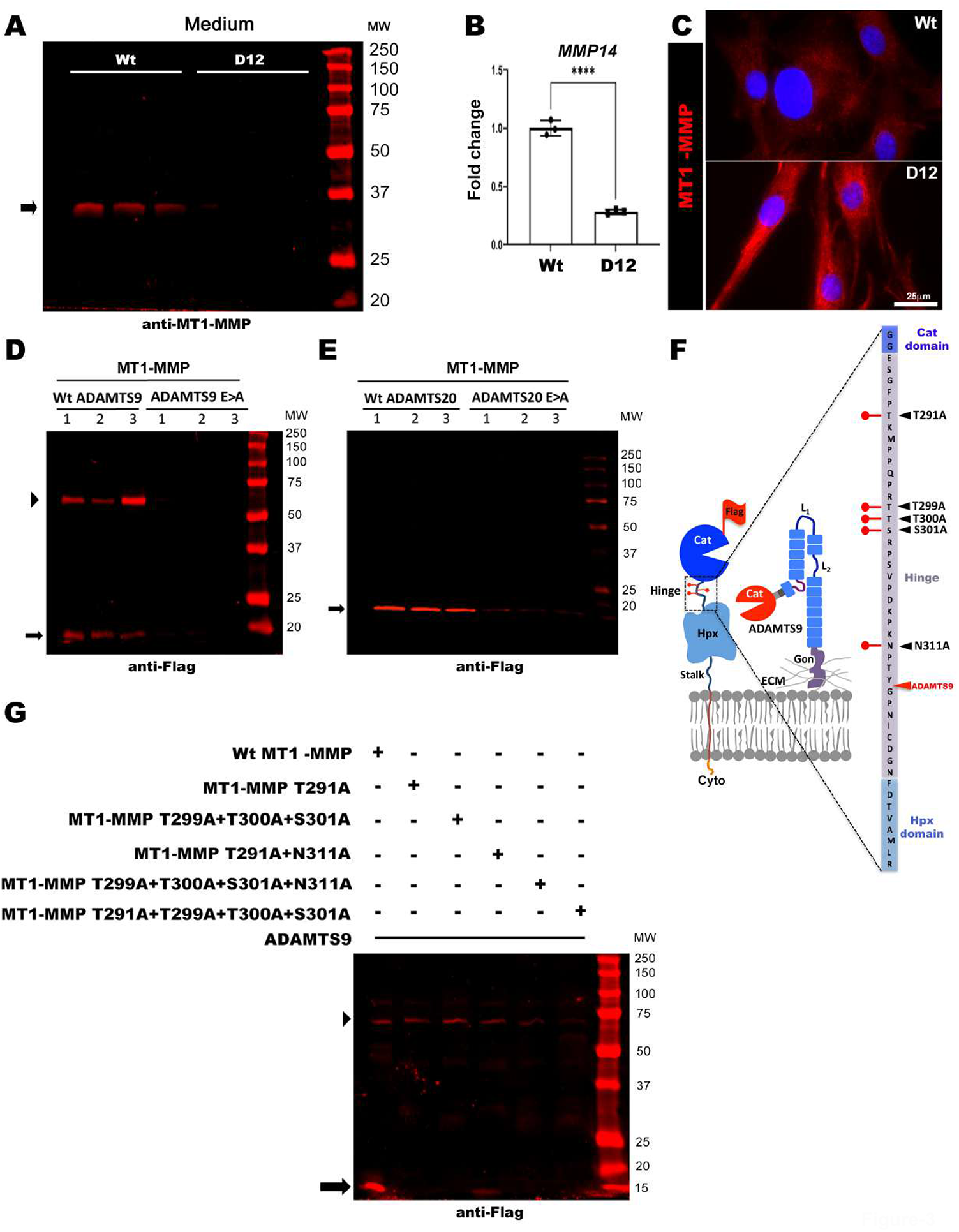
MT1-MMP is cleaved by ADAMTS9 and ADAMTS20 and MT1-MMP hinge glycosylation is essential for proteolysis by ADAMTS9. **(A)** Western blot of parental RPE-1 (Wt) and D12 conditioned medium with a catalytic domain-specific antibody showing release of the MT1-MMP catalytic domain (arrow indicates the catalytic domain fragment) in Wt, but not D12 cells. **(B)** Quantitative RT-PCR analysis of *MMP14* (MT1-MMP) transcript levels in parental RPE-1 (Wt) and D12 cells shows significantly downregulated expression in D12 cells. Error bars indicate S.D., **** p<0.0001, two-tailed unpaired Student t-test. **(C)** Cell surface immunostaining of MT1-MMP utilizing a catalytic domain antibody (red) shows more intense MT1-MMP staining in D12 cells. Nuclei are stained blue by DAPI. Molecular weight markers are shown on the right of the blot. Scale bar is 25μm. **(D)** Western blot (anti-FLAG) of the medium of HEK293T cells co-transfected in triplicate with MT1-MMP and ADAMTS9 showing molecular species of ∼60 kDa (arrowhead), corresponding to the MT1-MMP ectodomain and presumed to occur within the MT1-MMP stalk and ∼15 kDa (arrow), which are lacking in the medium of cells co-transfected with catalytically inactive ADAMTS9 (ADAMTS9E>A). Molecular weight markers are shown on the right of the blot adjoining the protein ladder. **(E)** Western blot (anti-FLAG) of the medium of HEK293T cells co-transfected in triplicate with MT1-MMP and ADAMTS20 showing a single 15 kDa fragment (arrow), which is lacking in the medium of cells co-transfected with catalytically inactive ADAMTS20 (ADAMTS20E>A). Molecular weight markers are shown on the right of the blot. **(F)** Schematic illustrating the MT1-MMP hinge in greater detail and depicting a model of ADAMTS9 cleavage at the MT1-MMP hinge. The hinge sequence is shown at far right flanked by the catalytic (Cat) domain and hemopexin (Hpx) domains. Lollipops indicate the glycosylated residues, and black arrowheads the specific mutations at these sites. The ADAMTS9 cleavage site is indicated by the red arrowhead. Gon, Gon-1 domain, ECM, extracellular matrix, Cyto, cytoplasmic tail. **(G)** Western blot of the medium from HEK293T cells co-transfected with ADAMTS9 and MT1-MMP or the indicated glycosylation mutants. Note the absence of the ∼15 kDa fragment indicated by the arrow) in all glycosylation mutants and additionally, lack of stalk cleavage in the T291A+T299A+T300A+T301A mutant (∼60 kDa fragment indicated by arrowhead). Molecular weight markers are shown on the right of the blot.

### *O*-glycosylation of the MT1-MMP hinge domain is required for ADAMTS9-mediated catalytic domain shedding

The MT1-MMP hinge contains potential sites for *O-*glycosylation (at Thr^291^, Thr^299^, Thr^300^ and Ser^301^) and one potential site for N-glycosylation at Asn^311^(Fig. 3F), although Pro at the Xaa position in the consensus N-glycosylation sequence, Asn-Xaa-Ser/Thr, suggests that this modification could be lacking (42). *O-*glycosylation of the MT1-MMP hinge protects it from autolysis (43), suggesting a potential influence on proteolysis by ADAMTS9 whereas its loss results in a closed catalytic domain pocket inaccessible for interaction with TIMP-2, which directly binds to and recruits pro-MMP2 for activation (44). We therefore inquired if MT1-MMP *O*-glycosylation was required for ADAMTS9-mediated proteolysis of MT1-MMP. Co-transfection of ADAMTS9 with wild type or mutant MT1-MMP with mutations of glycosylated sites (Thr^291^Ala, Thr^299^Ala, Thr^300^Ala, Ser^301^Ala, Asn^311^Ala) showed reduced proteolysis at the hinge, although cleavage at the presumed C-terminal site that generated the 60 kDa fragment was unaffected (Fig. 3G). Quadruple combined mutations of *O*-glycosylated sites (Thr^291^Ala + Thr^299^Ala + Thr^300^Ala + Ser^301^Ala + Asn^311^Ala) caused loss of both the ∼15 kDa and ∼60 kDa bands (Fig. 3G). These results suggest that *O*-glycosylation of the MT1-MMP hinge is necessary for ADAMTS9-mediated proteolysis of MT1-MMP.

### Degradomics identification of a second ADAMTS9 cleavage site in the MT1-MMP hemopexin domain

Because of the type 1 membrane topology of MT1-MMP, detection of an N-terminally labeled peptide in the conditioned medium that reported cleavage at Tyr^314^-Gly^315^ suggested occurrence of one or more downstream cleavages that released this peptide from the membrane anchored MT1-MMP. At least one such cleavage was evident from the 60 kDa fragment detected upon co-transfection of MT1-MMP and ADAMTS9. We investigated the possibility of additional cleavage sites both by targeted searches of the TAILS and pre-TAILS datasets from the medium for additional MT1-MMP peptides and 8-plex iTRAQ-TAILS comparison of RPE-1 and D12 cell lysate.

We first analyzed the medium TAILs and pre-TAILS data for differentially abundant MT1-MMP peptides with non-tryptic C-termini that could indicate a cleavage site and also had internal labeled lysines for quantification (Fig. 4A). This strategy identified two peptide spectrum matches for peptide, **GLPTDKIDAALFWMPNGKTYF**^429^↓(F^430^) (MS-identified sequence in bold letters, flanking amino acid in parentheses, with the residues at the non-tryptic site numbered) (Fig. 4B-D). This peptide is tryptic (preceded by Arg) at its N-terminus, and was not modified at the N-terminus by reductive dimethylation, but is non-tryptic at its C-terminus. It is located in the third blade of the four-bladed β-propeller structure of the hemopexin domain, which is bookended by two Cys residues that form the single disulfide bond in this domain. The peptide could be quantified from the dimethyl isotope labeling of two internal Lys residues, which also protected them from trypsin cleavage and was more abundant in wild-type cells (Fig. 4C,D). Cleavage at the F^429^-F^430^ site is unlikely to partition MT1-MMP into fragments, becaue of the disulfide bond between the first and fourth blades of the β-propeller although it could modify the tertiary structure of the hemopexin blade, β-propeller or full-length MT1-MMP (Fig. 4E).

**Figure 4.**
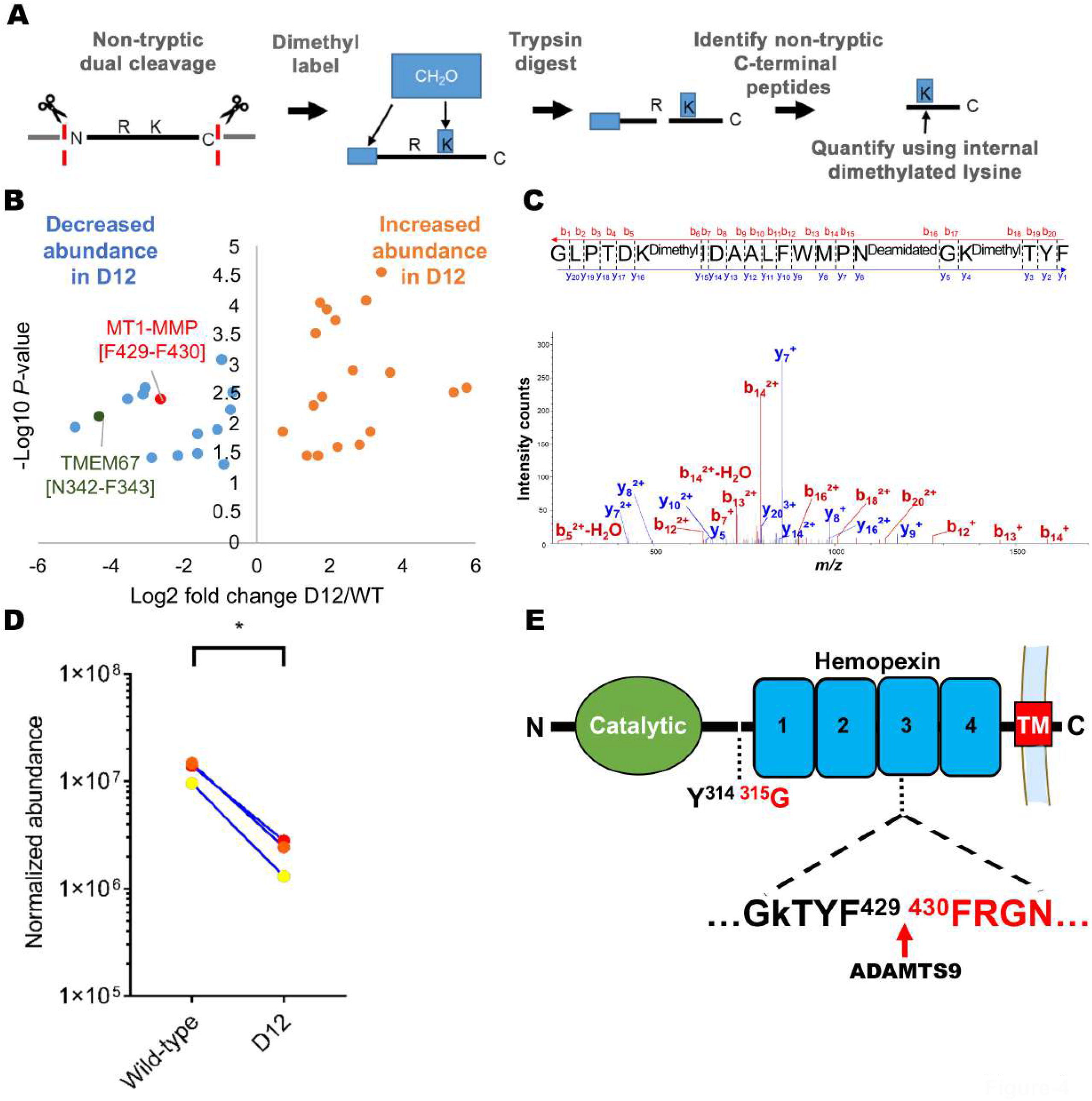
Degradomics identification of a cleavage site in the MT1-MMP hemopexin domain. **(A)** Schematic of the strategy for identifying non-tryptic peptides in the medium of parental RPE-1 cells and D12 cells and quantitation via internal dimethyl-labeled lysine residues. **(B)** Volcano plot of peptides identified with this strategy highlighting MT1-MMP and TMEM67 peptides and indicating cleavages at Phe^429^-Phe^430^ (red) and Asn^342^-Phe^343^ (dark blue), respectively. **(C)** MS2 profile of the MT1-MMP peptide GLPTDKIDAALFWMPNGKTYF. **(D)** Quantitation of normalized abundance of the identified MT1-MMP peptide in triplicate experiments. ** p<0.005, two-tailed unpaired Student t-test. **(E)** Location of the deduced cleavage site in blade 3 of the hemopexin domain. The hinge cleavage site is also shown. TM, transmembrane segment.

Next, among differentially abundant iTRAQ-labeled MT1-MMP peptides identified in RPE-1 and D12 cell lysates, the only informative differentially abundant peptide had the sequence (Thr^313^) ↓**Y^314^ GNICDGNFDTVAMLR** (MS-identified peptide sequence in bold letters, flanking amino acid in parentheses), suggesting cleavage at Thr^313^-Tyr^314^ (Supplemental Figure 1A-B). In contrast to the prior two identified peptides, this peptide was more abundant in the D12 cell lysate suggesting a cleavage that was suppressed in the presence of ADAMTS9 (Supplemental Figure 1C).

### ADAMTS9 knockout is associated with increased MMP2 zymogen activation and enhanced collagenase activity

Cell-surface MT1-MMP is well established as an activator of pro-MMP2 (33). Gelatin zymography using conditioned medium from parental RPE-1 cells and D12 cells showed increased pro-MMP2 activation (i.e., more of the 62kDa active MMP-2, as well as a 65 kDa intermediate form) in the absence of ADAMTS9 (Fig. 5 A,B). Culture of parental RPE-1 and D12 cells on DQ-collagen I coated slides revealed greater collagenase activity under D12 cells, which was revealed by the fluorescence de-quenching resulting from substrate proteolysis (Fig. 5C). These results suggest that ADAMTS9 could potentially regulate MT1-MMP activity on the cell-surface directly in addition to downstream activation of pro-MMP2. Intriguingly, increased MMP2 activity noted on zymograms was not a consequence of higher *MMP2* mRNA, which in fact was significantly lower in the D12 cells (Fig. 5D).

**Figure 5.**
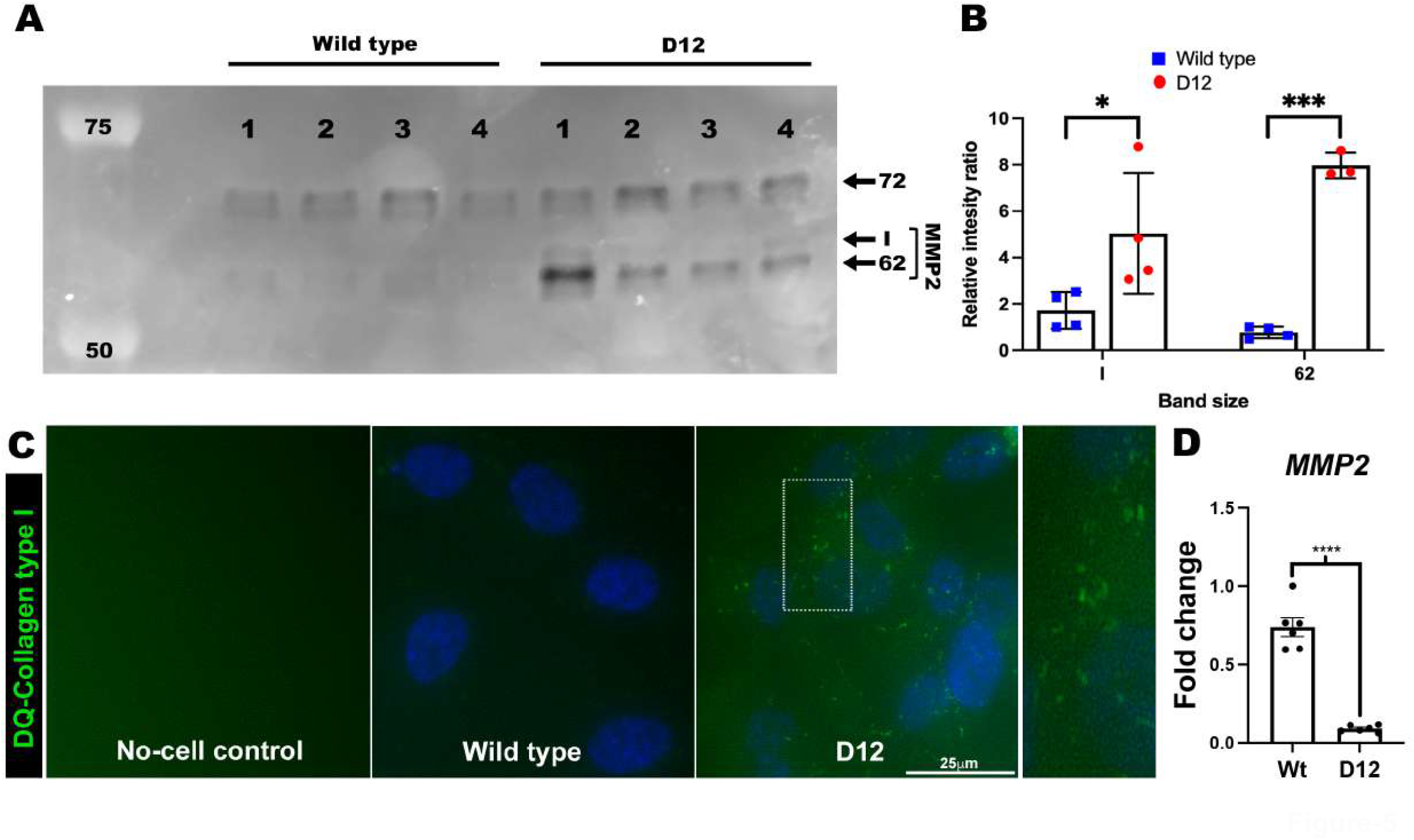
Increased pro-MMP2 activation and collagenase activity in D12 cells as a result of increased cell-surface MT1-MMP. **(A)** Gelatin zymography of the medium from parental RPE-1 cells (wild-type) and D12 cells shows increased active MMP2 (62 kDa) as well as an intermediate species (I) in the D12 medium. **(B)** Quantitation of band intensities from A show significantly higher active MMP2 levels in the D12 medium. N=4, each group, error bars indicate S.D., * indicates p<0.05, *** indicates p<0.05, Student t-test. **(C)** Parental RPE-1 cells and D12 cells were cultured on DQ collagen type I coated plates for 24 hours and areas of collagenolysis were visualized by green fluorescence, showing greater activity in D12 cells compared to parental RPE-1 cells. **(D)** Quantitative RT-PCR analysis shows reduced *MMP2* mRNA in D12 cells. Error bars indicate S.D., ****, p<0.0001, two-tailed unpaired Student t-test. Scale bar in C is 25μm.

### MT1-MMP modulates cilium length in RPE-1 cells, but without affecting cilium biogenesis

ADAMTS9 is required for primary cilium biogenesis in humans and in mice, i.e., the primary cilium either fails to form in cells of *Adamts9* or *Adamts9+Adamts20* mutant mice in vivo and in D12 cells after serum starvation (primary cilia form in post-mitotic cells), or if formed, is extremely short (12, 25). We therefore inquired if alteration of cell-surface MT1-MMP levels affected ciliogenesis. First, using an siRNA which provided substantial MT1-MMP depletion, (Fig. 6A,B), we observed no impact on the percentage of parental RPE-1 cells having primary cilia after serum starvation although cilium length was slightly increased compared to cells transfected with control siRNA (Fig. 6C,D). Overexpression of MT1-MMP in parental RPE-1 cells did not alter the percentage of ciliated cells either but resulted in slightly shorter primary cilia compared to cells transfected with the empty vector (Fig. 6C,D). Thus, in cells with ADAMTS9 expression, increasing or decreasing MT1-MMP had no impact on ciliogenesis, although a minor effect on cilium length was observed. To inquire if loss of cilia in the absence of ADAMTS9 could be explained by increased MT1-MMP activity, we depleted MT1-MMP in D12 cells and induced ciliogenesis by serum-starvation. Although MT1-MMP knockdown led to even lower levels than in parental RPE-1 cells as judged by immunofluorescence of unpermeabilized cells (lower panels, Fig 6A), neither the percentage of ciliated cells nor cilium length were affected (Fig. 6E,F). These results indicate loss of cilium biogenesis observed in the absence of ADAMTS9 was not due to increased cell-surface MT1-MMP or greater MMP2 activation.

**Figure 6.**
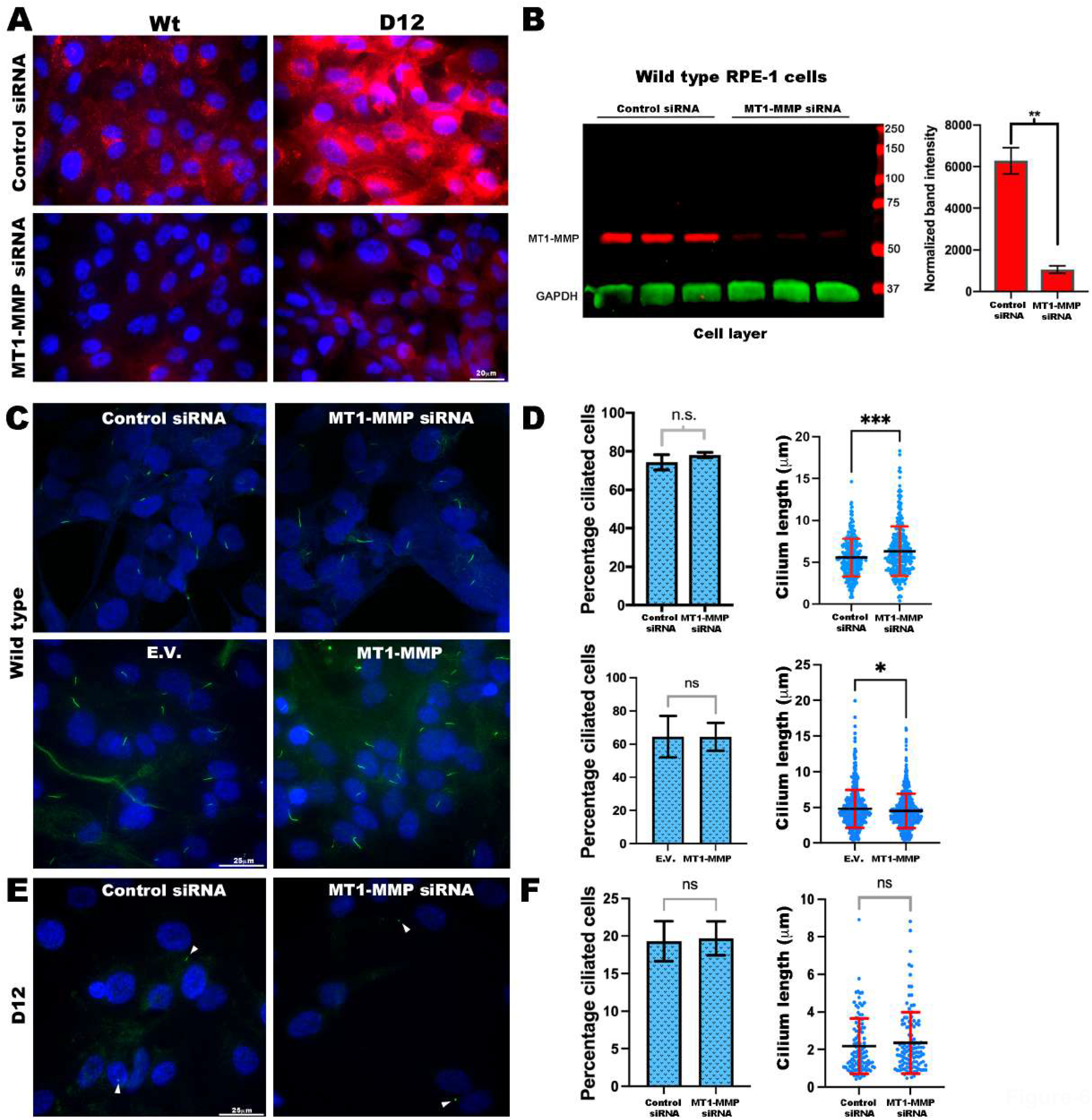
Depletion or over-expression of cellular MT1-MMP affects cilium length, but not cilium biogenesis. **(A)** Immunostaining of cell surface MT1-MMP (red) shows that *MMP14* siRNA effectively depleted MT1-MMP in both parental RPE-1 cells (Wt) and ADAMTS9-edited RPE-1 D12 cells. **(B)** Western blot of parental RPE-1 cells treated with control siRNA or *MMP14* siRNA shows significant depletion of MT1-MMP (red) normalized to GAPDH (green) as a measure of knockdown. Error bars indicate S.D., ** indicates p<0.005, two-tailed unpaired Student t-test analysis. (**C-D**) Primary cilium staining using acetylated-α-tubulin antibody (green) in parental RPE-1 cells treated with control siRNA and *MMP14* siRNA (upper panels) or transfected with MT1-MMP or empty vector (E.V.) (lower panels), showed that MT1-MMP depletion did not affect the percentage of ciliated cells but slightly increased cilium length. MT1-MMP over-expression also did not affect the percentage of ciliated cells but slightly decreased cilium length. Red error bars indicate S.D., black lines indicate mean, ***, p<0.0005, *, p<0.05 two-tailed unpaired Student t-test. (**E-F**) Primary cilium analysis using acetylated-α-tubulin immunostaining (green) after depletion of MT1-MMP in D12 cells showed no impact on the percentage of ciliated cells nor cilium length. Red error bars indicate S.D., black lines indicate the mean. Scale bars are 20 μm in A and 25 μm in B.

### Reversal of reduced adhesion in ADAMTS9-deficient D12 cells by MT1-MMP knockdown

To inquire if impaired formation of focal contacts in D12 cells (Fig 1A) was due to increased MT1-MMP activity, we repeated IRM after MT1-MMP knockdown (Fig 7A). The time-lapse IRM imaging demonstrated accelerated emergence of focal adhesions 30 min after seeding D12 cells treated with *MMP14* siRNA compared to control siRNA treatment (Fig. 7B). In these cells, *MMP14* siRNA was clearly effective in reducing its expression, whereas there was no impact on *MMP2* expression (Fig. 7C). 24 h post seeding, both parental RPE-1 and D12 cells with *MMP14* knockdown demonstrated a shift in the relative proportion of close contacts (gray) and focal/fibrillar adhesions (dark areas), with increased focal adhesions evident in the D12 cells in response to MT1-MMP depletion (Fig. 8A,B). Intriguingly, the trailing edge membrane protrusions observed in D12 cells were not observed after MT1-MMP depletion (Movies S3-4). Quantification of binned pixel intensities in the spectrum of dark to bright pixels per cell demonstrated a statistically significant gain of dark pixels in D12 cells in response to MT1-MMP depletion, whereas parental RPE-1 cells exhibited a statistically significant reduction of the bright pixels and a moderate increase of the darker pixels, albeit statistically insignificant (Fig. 8C).

**Figure 7.**
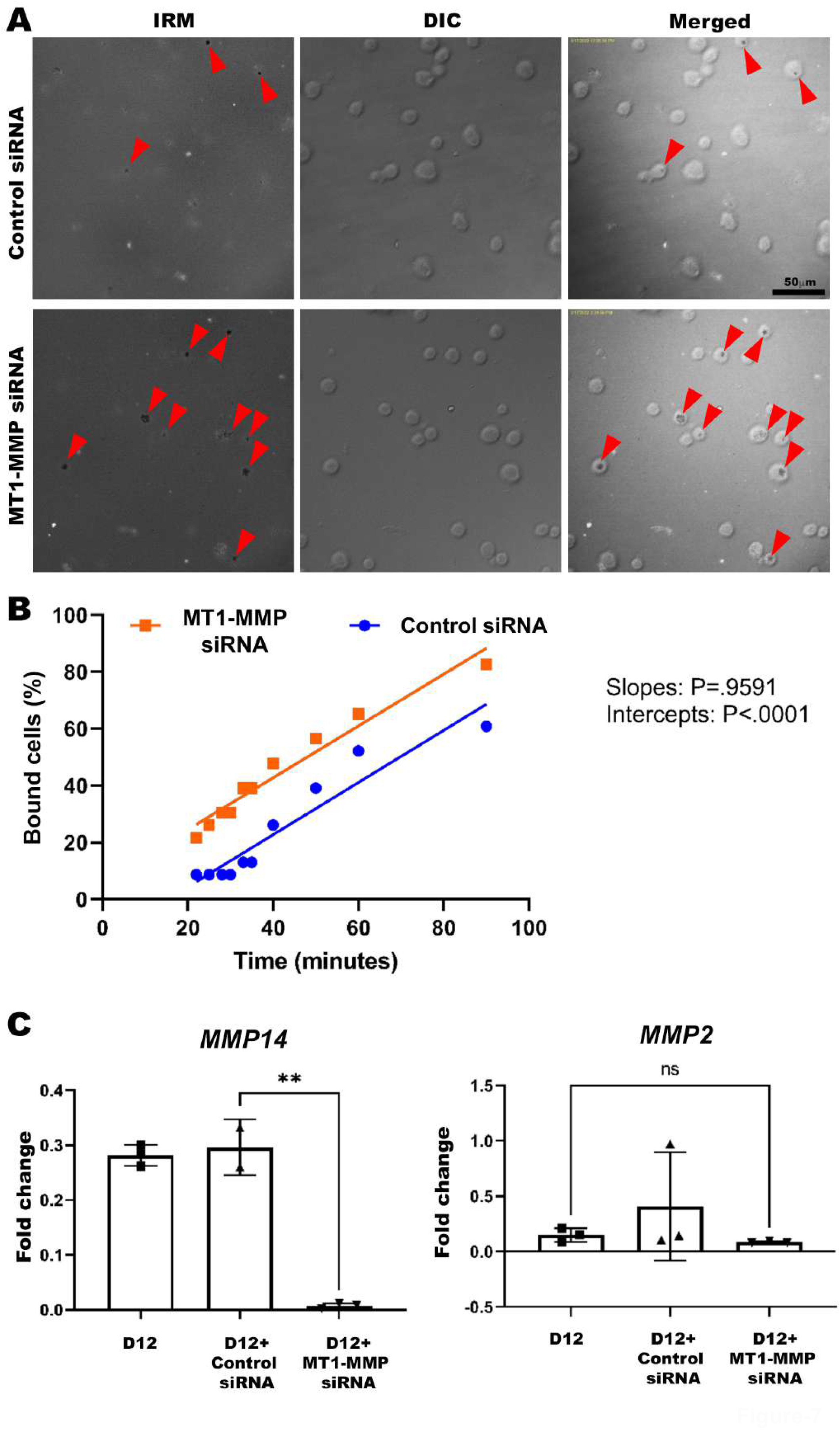
MT1-MMP depletion restored cell-substratum interaction in RPE1-ADAMTS9^KO^ cells. **(A)** Still images taken from concurrent, aligned interference reflection microscopy (IRM) and differential interference contrast (DIC) time-lapse imaging show reversion of the cell-substratum interface to that noted at the same time point in parental RPE-1 cells (see Fig. 1A) after *MMP14* knockdown. Red arrowheads indicate attached cells with dark contacts. **(B)** Percentage of D12 cells bound by focal adhesions observed by time-lapse microscopy after transfection with MT1-MMP siRNA (orange) and scrambled siRNA (blue) shows consistently better cell attachment in the former. **(C)** Quantitative RT-PCR analysis for *MMP14* and *MMP2* transcripts in D12 cells treated with control siRNA and *MMP14* siRNA shows effective knockdown and suggests that stronger attachment after *MMP14* knockdown is not due to altered transcription of MMP2. Error bars indicate S.D., ** indicates p<0.0005 in two-tailed unpaired Student t-test. Scale bar is 50μm in A.

**Figure 8.**
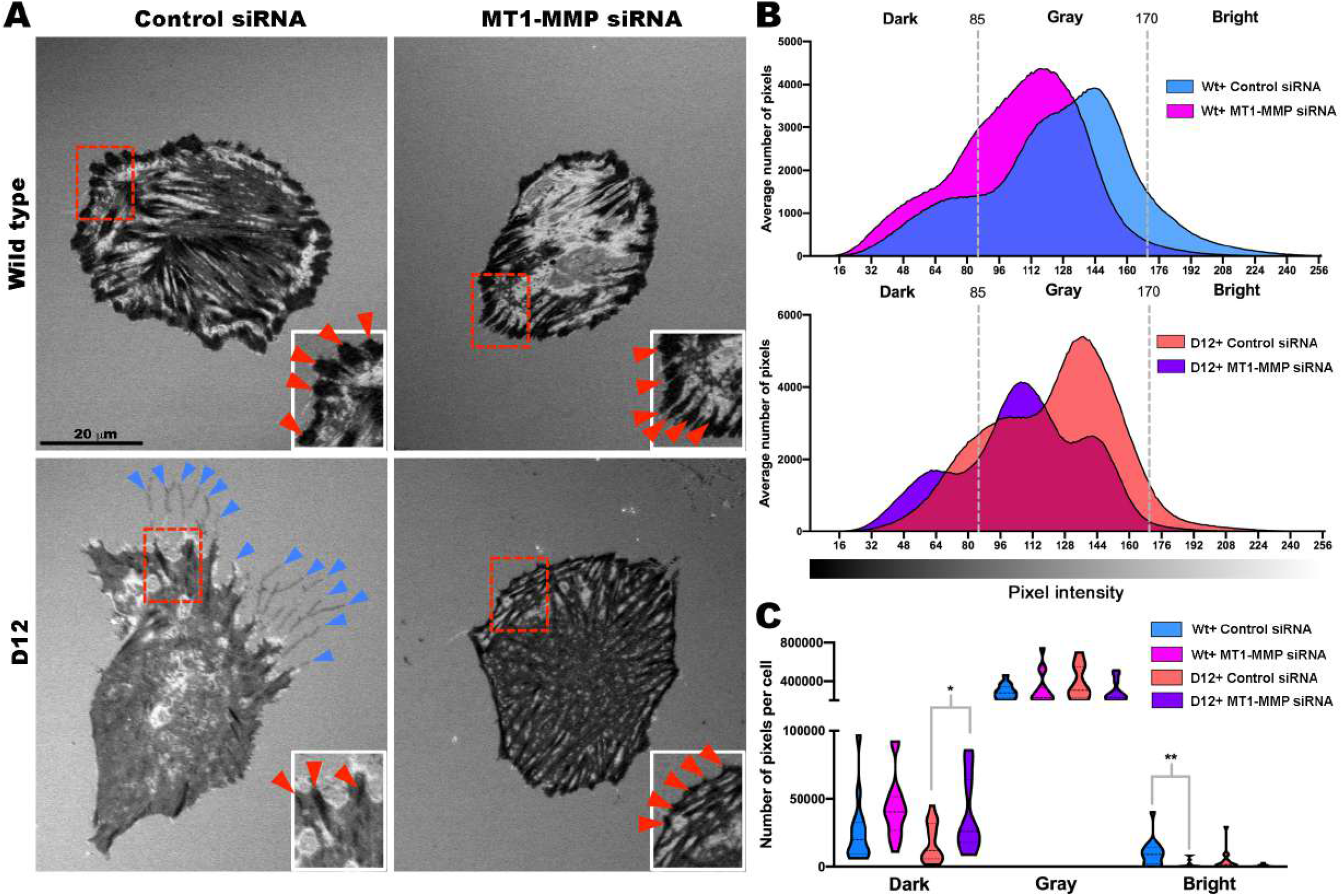
MT1-MMP knockdown improves focal adhesion formation in D12 cells. **(A)** Interference reflection microscopy (IRM) imaging 24 hours after seeding of parental RPE-1 cells and D12 cells treated with control siRNA or *MMP14* siRNA shows an increase in fibrillar adhesions, especially at the cell periphery (red arrowheads, inset box) and lack of trailing edge filamentous extensions observed in D12 cells (blue arrowheads). **(B)** Distribution of IRM pixel intensities comparing parental RPE-1 cells treated with control siRNA (blue) or MT1-MMP siRNA (pink) and D12 cells treated with control siRNA (salmon) or MT1-MMP siRNA (purple). Dashed lines indicate binning of dark (0-85), gray (85-170) and bright (170-256) pixels. **(C)** Violin plot of binned pixel intensities per cell showing increased dark pixels in D12 cells treated with MT1-MMP siRNA compared to control siRNA treatment and decreased bright pixels in wild type cells treated with MT1-MMP siRNA. N=14-16 cells per treatment group, * p<0.05, ** p<0.005, two-tailed unpaired Student t-test. Scale bar=20μm in **A**.

## Discussion

The present work shows that RPE-1 cells lacking ADAMTS9 delay initiation of focal adhesions and form fewer peripheral fibrillar adhesions with a glass substratum than parental RPE-1 cells. Thus, a lack of ADAMTS9 led to dramatic alterations in their initial attachment and spreading. Our degradomic analysis of these cells, which was undertaken to define the proteolytic changes possibly associated with this effect, identified potential substrates requiring further investigation. Here, we undertook detailed validation and further analysis of MT1-MMP as a substrate, since this cell-surface protease is a major participant in ECM proteolysis, both directly and via activation of other proteases (45, 46), and like ADAMTS9 and ADAMTS20, is strongly implicated in embryonic development (47-50), albeit without any correspondence of the respective phenotypes. In contrast to *Adamts9*, whose functions span several developmental processes from gastrulation to birth, the phenotypes of *Mmp14-*deficient mice primarily reflect its involvement in fibrillar collagen proteolysis in the postnatal and juvenile period (47, 51). This lack of correspondence between the phenotypes of *Adamts9* and *Mmp14* deficient mice does not detract from the significance of the present findings, since our data suggest that a lack of ADAMTS9 and ADAMTS20 leads to increased cell-surface MT1-MMP, for which no genetic model presently exists. *Adamts9* null mice do not survive past gastrulation (15), and *Adamts9* and *Adamts20* mRNAs are co-expressed in many tissues, so that it may be challenging to determine whether MT1-MMP activity is specifically increased in embryos in the absence of ADAMTS9 and ADAMTS20.

Most endopeptidases have selective, albeit broad, but not indiscriminate substrate repertoire, and it is only the rare protease which has a private substrate (e.g., ADAMTS13 appears to cleave only von Willebrand factor). The first identified ADAMTS9 substrates were the aggregating ECM proteoglycans aggrecan and versican (8). TAILS analysis of skin from 1 week-old wild-type and an *Adamts9* mutant (*Adamts9*^Und4/+^ (31)) identified several ECM components as potential substrates, although none were orthogonally validated. The yeast two-hybrid system was used to screen for ADAMTS9 binding partners, identifying the ECM glycoprotein fibronectin as a potential substrate (52). Proteomics analysis of fibronectin semi-tryptic peptides in the presence of catalytically active ADAMTS9 identified a new cleavage site and ADAMTS9 proteolytic activity was shown to impair fibronectin fibril assembly by cultured fibroblasts (52). The present work, using TAILS and a different cellular system, has identified additional potential fibronectin cleavages, although not the previously identified site. Thus, ADAMTS9 cleaves three orthogonally validated ECM substrates and likely others, such that a tissue or cell phenotype resulting from ADAMTS9 deficiency could have a complex mechanistic basis reflecting accumulation of multiple substrates.

The impaired initial adhesion and observed reduction in fibrillar adhesions in D12 cells is consistent with prior work showing that cultured human myometrial smooth muscle cells (SMCs) did not form fibrillar adhesions and were poorly adherent after *ADAMTS9* knockdown (14). Myometrial SMCs, maintained from primary cultures, appear to depend heavily on focal adhesions for their survival, since they subsequently underwent anoikis after *ADAMTS9* knockdown (14). However, the CRISPR-Cas9 generated *ADAMTS9*-mutant D12 clonal derivative of RPE-1 cells (an immortalized cell line), could be maintained in culture despite a complete lack of ADAMTS9 (12). Versican had previously emerged as a key anti-adhesive ADAMTS9 substrate in human myometrial SMCs, since they formed few focal adhesions after *ADAMTS9* knockdown and versican accumulated in *Adamts9*-deficient mouse myometrium; moreover, focal adhesions were restored dramatically by versican knockdown concomitant with ADAMTS9 knockdown in human myometrial SMCs (14). Versican, which also accumulates in the ECM of D12 cells when cultured for 72 h or longer (12, 34), is likely also relevant to altered adhesion of D12 cells, although the possibility was not specifically experimentally addressed here. MT1-MMP knockdown had a restorative effect on fibrillar adhesion formation by D12 cells, suggesting that the complex tissue and cell phenotypes resulting from ADAMTS9 and ADAMTS20 deficiency reflect the impaired cleavage of more than one substrate, i.e., MT1-MMP, versican and potentially others.

Sequence analysis of cleavage sites identified in comparative TAILS of wild-type and *Adamts9*^Und4/+^ skin suggested a strong preference for aromatic residues Tyr and Phe or the hydrophobic residue Leu at the P1 position (according to the nomenclature of Schecter and Berger) and Thr at P2. The two cleavage sites attributed to ADAMTS9 in MT1-MMP are consistent with this preference, with Tyr and Phe at the P1 position and Gly or Phe, respectively, at P1’
s. The processing event in the MT1-MMP hinge with higher abundance in D12 cells (Thr^313^-Tyr^314^) may be a constitutive cleavage, such as by autolysis, that may occur if prior processing by ADAMTS9 is lacking. MT1-MMP autolysis was previously reported to occur at Gly^284^-Gly^285^ in the hinge, followed by subsequent cleavage at Ala^255^-Ile^256^ (53, 54). Paradoxically, it was reported that the residual cell-surface attached ectodomain is associated with increased cell surface proteolytic activity, because it impairs endosomal intake of full-length MT1-MMP, resulting in higher levels of active enzyme at the cell surface, which may in part explain increased MT1-MMP staining and activity we noted despite reduced *MMP14* transcription (55). The regulation of cell-surface MT1-MMP by ADAMTS9/20 adds to the complex regulatory mechanisms of MT1-MMP which were previously reviewed (33, 56).

Prior publications have shown that MT1-MMP regulates focal adhesions and the findings of the present study are consistent with the previously observed effects. MT1-MMP was previously shown to attenuate integrin clustering in HT1080 cells adherent to fibronectin; in contrast, MT1-MMP inhibition led to formation of stable adhesions (57, 58). It has also been shown in several cell lines that MT1-MMP mediates proteolysis of fibronectin and other substrates at focal adhesions (59), including ARPE-19 cells, which like the RPE-1 cells used in this study, are derived from retinal pigment epithelium. Prior work suggested that MT1-MMP is localized to focal adhesions via interaction with β1-integrin (60), but it remains to be established whether ADAMTS9 or ADAMTS20 interact directly with integrins at the cell-surface and whether they proteolytically cleave MT1-MMP at focal adhesions, elsewhere on the cell surface, in the secretory pathway or in endosomes. Woskowicz et al have shown that the MT-loop, a segment of the catalytic domain of MT1-MMP ((163)PYAYIREG(170)), i.e, likely within the putative N-terminal regions released by ADAMTS9/20 from the cell-surface, is essential for MT1-MMP promotion of cellular invasion by localization in β1-integrin-rich cell adhesion complexes at the plasma membrane, whereas other work has suggested a key role for the cytoplasmic tail and transmembrane domain in β1-integrin regulation (61). MT1-MMP was also shown to co-localize and interact with α_v_β_3_ integrin, which it proteolytically cleaves (62-64). Notably, MT1-MMP-deficient myeloid progenitors and mouse lung endothelial cells have larger focal adhesions (65). It was postulated that this effect was potentially related to accumulation of adhesive proteins such as fibronectin and collagen I which are MT1-MMP substrates and a lower turnover of focal adhesions (66). Our data, showing that lack of ADAMTS9/20 led to fewer focal adhesions in RPE1 cells, with greater cell-surface collagenase activity and MMP2 activation, appear to be consistent with the expected effect of increased MT1-MMP at the cell surface.

Previously, enzymatic deglycosylation, site-directed mutagenesis, and lectin precipitation assays were used to demonstrate that the MT1-MMP hinge region contains *O-*linked carbohydrates on Thr^291^, Thr^299^, Thr^300^, and/or Ser^301^(44). MT1-MMP autolysis is increased in the absence of glycosylation (43) and MT1-MMP lacking glycosylation at these sites is unable to bind TIMP-2, preventing formation of the MT1-MMP/TIMP2/proMMP2 trimeric complex essential for pro-MMP2 activation (44). The positive effect of MT1-MMP hinge glycosylation on susceptibility to cleavage by ADAMTS9 extends prior work identifying an influential role for N-linked and *O*-glycosylation on the activity of MT1-MMP and other proteases (67-70).

*MMP21* mutations can cause heterotaxy (71-73), a ciliopathy, but there is no evidence to date that MT1-MMP is required for the formation or function of cilia, since mouse MT1-MMP mutants lack the morphogenetic defects that are typically associated with primary cilium dysfunction. Although the primary cilium is distinct from motile cilia present on many epithelial cells, *Mmp14* knockout mice do however have defective ependymal motile cilia, which are thought to underlie hydrocephalus in these mice (74). The primary cilium was not previously examined in MT1-MMP deficient states, and our data showed that MT1-MMP knockdown has no impact on cilium biogenesis, although cilia are slightly longer than in parental RPE-1 cells. Although lack of ADAMTS9 in D12 cells leads to increased cell-surface MT1-MMP, and not its inactivation, observation of motile cilium defects in MT1-MMP mutant mice is nevertheless intriguing since hydrocephalus is also observed in *Adamts20*^Bt/Bt^ mice (75), although its mechanism is yet to be resolved.

The present findings are conceptually broadly significant for the protease web, in which positive and negative feedback loops, including activation or elimination of protease activity, are mediated by proteases within the same or different families (76). For example, MT1-MMP activates MMP-2 and MMP-13 in a feed-forward mechanism, whereas it inactivates the transmembrane protease ADAM9 (77). Here we expand the interactions of the MT1-MMP node in the web by establishing that ADAMTS9 and ADAMTS20 release the MT1-MMP catalytic domain to moderate its cell-surface activity (reported in the present study using MMP2 activation and collagen breakdown as readouts). In contrast to the effect of the transmembrane metalloprotease ADAM12 in activating MT1-MMP via a non-proteolytic mechanism that promotes pro-MMP2 activation (78), ADAMTS9 and ADAMTS20 are shown here to cleave MT1-MMP and suppress proMMP-2 activation as a potential physiological role. Although the findings of this study were elucidated exclusively in RPE-1 cells, it is possible that ADAMTS9 and ADAMTS20 may act similarly in tissues where they are expressed alongside MT1-MMP, such as during ocular development and in vascular endothelium or craniofacial mesenchyme.

ADAMTS9 is constitutively expressed in endothelial cells in most mouse capillary beds and is anti-angiogenic (10), which could be potentially explained by modulation of cell-surface MT1-MMP and MMP2 activity, which are strongly pro-angiogenic (79) and focal adhesion formation. In this context, genome-wide association studies in humans have associated the *ADAMTS9* locus with age-related macular degeneration (AMD) (80, 81), a major cause of adult-onset blindness, where RPE cell death and dysregulated angiogenesis are prominent features. In eyes at high risk of AMD, one study identified lower *ADAMTS9* mRNA levels (82), which we speculate could lead to higher MT1-MMP activity, and thus greater pro-MMP2 activation, together promoting retinal angiogenesis. In future work, it will be important to apply the findings disclosed here to mechanisms of eye disease, since ADAMTS9 is essential for eye development and is expressed along with MT1-MMP in the drainage apparatus, where it could have a role in intraocular pressure regulation (17, 83). In addition to fibronectin, previously identified as an ADAMTS9 substrate and other ECM and secreted proteins and proteoglycans, the present work also identified the cilium transition zone protein TMEM67, which like ADAMTS9, is mutated in ciliopathies as a potential substrate, warranting further investigation of TMEM67 processing in the context of ciliogenesis (84-86).

## METHODS

### Cell culture, recombinant DNA and siRNA transfections

Wild type and *ADAMTS9* KO (D12) RPE-1 cells (ATCC, CRL-4000) were cultured in DMEM F/12 culture medium containing 10% fetal bovine serum (FBS), 50 U/ml penicillin/streptomycin and 200 μg/ml Hygromycin B (Millipore Sigma, Cat. # 400052) as previously described (12). HEK293T cells (ATCC, CRL-1573) were cultured in DMEM with 10% FBS and 50U/ml penicillin/streptomycin. Plasmids encoding full length human ADAMTS9 and mouse ADAMTS20 and their catalytically inactive mutants (E/A) in pCDNA3.1 Myc/His vector as previously described (12). Wild type or catalytically inactive human MT1-MMP constructs with an N-terminal Flag tag between the propeptide and catalytic domain) were previously described (87). Human MT1-MMP hinge *N-* and *O*-glycosylation mutants T299A (pCR-CHO-3/f), T299A/T300A/S301A (pCR-CHO-3/f), T291A/N311A (pCR-NACHO-1/f), T299A/T300A/S301A/N311A (pCR-NACHO-3/f) and T291A/T299A/T300A/S301A (pCR-NACHO-4/f) were previously described (44). All plasmid DNA transfections were carried out using the PEI Max transfection reagent. siRNA targeting human MT1-MMP (Ambion, catalog no. AM51331) or a Negative Control (Invitrogen catalog no. 4390843) was transfected to wild type or D12 RPE-1 cells cultured in 8-chamber slides as previously described (Nandadasa et al., 2019) using the Lipofectamine RNAiMAX reagent (Thermo Fischer Scientific Cat# 13778100).

### Western blotting and gelatin zymography

1 ml of serum-free conditioned medium from RPE-1 or HEK293T cells cultured in 6-well plates for 48h post-transfection was collected and the cell layers were harvested using 300 μl PBS containing 1% Triton X-100 and protease inhibitor complex. 6X Laemmli sample buffer containing β-mercaptoethanol was used to load 50 μl of each sample for western blot analysis using 10% SDS PAGE. For western blots, anti-FLAG polyclonal antibody (Sigma, 1:1000 dilution) and anti-human MT1-MMP rabbit monoclonal antibody EP1264Y against the catalytic domain (Abcam, catalog no. ab51074, 1:1000 dilution) were used followed by anti-rabbit secondary antibodies (Li-COR) at 1:15000 dilution and the membranes were imaged using a Li-COR Odyssey CLx infrared imager (Li-COR).

For gelatin zymography, 1 ml of serum-free conditioned medium from parental RPE-1 or D12 cells (n=4 cultures for both) was collected after 48 hours culture in a 12-well plate and EDTA-free protease inhibitor (Roche, catalog no.11836170001) was added. Medium was concentrated to ∼200 μl using a 10 kDa molecular weight cut-off spin column (Amicon, catalog no. UFC501096) and 0.3 μg of total protein was loaded in each well using 5X Laemmli sample buffer without β-mercaptoethanol. Gelatin zymography was carried out using 10% SDS-polyacrylamide gels containing 1% gelatin (Novex, ThermoFisher, catalog no. ZY00102BOX) following the manufacturer protocol (Invitrogen, ThermoFisher). Gels were run until the 37 kDa marker reached the bottom third of the gel and were immediately washed with water prior to incubation at room temperature in 1X renaturing buffer (Novex, ThermoFisher, LC2670) for 30 minutes. The gel was equilibrated in 1X developing buffer (Novex, ThermoFisher, LC2671) at room temperature for 30 minutes followed by addition of fresh developing buffer and overnight incubation at 37°C. Following a 10 min wash with water, the gels were stained for 1 hour with GelCode Blue (Thermo Fischer, catalog no.24590) and destained for at least 2 hours with distilled water. Digestion bands were imaged with the LiCOR Odyssey Clx imager and quantified using ImageJ densitometry software (NIH, Bethesda, MD).

### TAILS sample preparation and mass spectrometry

Proteins from lysates and phenol red-free, serum-free conditioned medium of parental RPE-1 and D12 cells were analyzed by TAILS using N-terminal labeling with iTRAQ or reductive dimethylation respectively. For making lysates, cells were scraped from 10 cm dishes at confluence, washed with PBS and pelleted by centrifugation for 5 minutes. Cells were lysed by addition of 2.35 ml of 0.05% SDS, 10 mM DTT in 100 mM HEPES, 10 mM EDTA, pH 8 with protease inhibitors (cOmplete mini, Millipore Sigma) to the pellet, followed by tip probe sonication for 80 seconds at a 20% amplitude using a Q-500 sonicator (Qsonica). DNA and RNA were degraded using 2 μl of benzonase (EMD Millipore Corp., catalog no. 70746-10KUN) at room temperature for 20 minutes and acetone-methanol precipitation was performed to precipitate proteins. This pellet was resuspended in 2.5 M GuHCl and 250 mM HEPES. Protein quantity was determined using a 592 nm bisinchonic acid assay (Thermo Scientific, catalog no. 23225) assay and normalized to 250 μg per channel. Samples were reduced and alkylated using 10 mM tris(2-carboxyethyl) phosphine (TCEP) and 25 mM iodoacetamide (IAA), respectively. Each sample was labeled with 2 units of each channel by adding 1 sample volume of dimethyl sulfoxide (DMSO) to each iTRAQ label (Sciex, catalog no.4390812) before adding the total solution volume to each sample. Four iTRAQ channels were used to label parental RPE-1 cell lysates and four were assigned to label D12 lysates for 2 hours in the dark at room temperature before quenching the labeling reaction with ammonium bicarbonate to a final concentration of 100 mM. The samples labeled with the respective iTRAQ labels were combined and proteins were precipitated using an acetone-methanol precipitation method.

The medium was passed through a 10 µm filter to remove cell debris and concentrated ten-fold using a 3kDa molecular weight cut-off (MWCO) spin filter (Amicon, catalog no. UFC500396). Protein was quantified as above and normalized to 500 μg per sample. Each sample was reduced and alkylated using 5 mM dithiothreitol (DTT) and 20 mM IAA, respectively, and excess IAA was quenched using an additional 15 mM DTT. Samples were labeled with 40 mM isotopically distinct dimethyl formaldehyde (light, CH_2_O, for the parental RPE-1 cells (Cambridge Isotope Laboratories, Inc., catalog no.ULM-9498-100) and heavy, ^18^CD_2_O for D12 (Sigma Aldrich, catalog no. 596388-1G) added simultaneously with 20 mM NaBH_3_CN and incubated overnight at 37°C. Fresh formaldehyde and NaBH_3_CN were added the following day for 1 h. The reaction was quenched in a final concentration of 100 mM Tris HCl at 37° C for 2 h. Paired heavy and light-labeled replicate samples were then mixed and the proteins separated by chloroform-methanol precipitation.

Precipitated pellets were solubilized in 100 mM NaOH and 500 mM HEPES pH 7.5. Trypsin was added at a 1:100 (protease: protein) ratio and incubated at 37° C overnight. A 30 μg aliquot was removed from each sample (the pre-TAILS sample) and the remaining protein was mixed with hyperbranched polyglycerol-aldehydes (HPG-ALD, Flintbox, https://www.flintbox.com/public/project/1948/) at a 5:1 polymer: protein ratio overnight at 37° C in the presence of 10 mM NaBH_3_CN. HPG-ALD binds unblocked (i.e. trypsin-generated) amino acid termini, excluding them from the sample and enriches for peptides with blocked/labeled N-termini (TAILS peptides). The samples were filtered through a 10 kDa MWCO column (EMD Millipore) and the eluate and pre-TAILS fractions were desalted on a C18 Sep-Pak column (Waters, catalog no. WAT054955) and eluted in 60:40 ACN: 1% TFA. Samples were vacuum-centrifuged until dry and resuspended in 1% acetic acid for mass spectrometry.

The pre-TAILS and TAILS peptides were analyzed on a ThermoFisher Scientific Fusion Lumos tribrid mass spectrometer system interfaced with a Thermo Ultimate 3000 nano-UHPLC. The HPLC column was a Dionex 15 cm x 75 µm id Acclaim Pepmap C18, 2 µm, 100 Å reversed phase capillary chromatography column. 5 µL volumes of the trypsin-digested extract were injected, peptides were eluted from the column by an acetonitrile/ 0.1% formic acid gradient at a flow rate of 0.3 µL/min and introduced in-line into the mass spectrometer over a 120-minute gradient. The nanospray ion source was operated at 1.9 kV. The digest was analyzed using a data-dependent method with 35% collision-induced dissociation fragmentation (for the dimethyl labeled samples) or high-energy collisional dissociation set to 38 (for the iTRAQ labeled) of the most abundant peptides every 3s and an isolation window of 0.7 m/z for ion-trap MS/MS. Scans were conducted at a maximum resolution of 120,000 for full MS. Dynamic exclusion was enabled with a repeat count of 1 and ions within 10 ppm of the fragmented mass were excluded for 60 seconds.

### Proteomics data analysis

Spectra were searched against the reviewed human database (November 2018) using Proteome Discoverer V2.2 Software (Thermo Fisher). Peptides were identified using a precursor mass tolerance of 10 ppm, and fragment mass tolerance of 0.6 Da with the static modification being carbamidomethyl (C), whereas dynamic modifications included either the light (28.03 Da) or heavy (32.06) dimethyl formaldehyde (N-terminal, K) or iTRAQ 8-plex (N-terminal, K) as well as oxidation (M), deamidation (N), acetylation (N-terminal), and Gln to pyro-Glu N-terminal cyclization. Peptides were validated using a false discovery rate (FDR) of 1% against a decoy database. Only high confidence proteins (containing peptides at a 99% confidence level or higher) were recorded from each sample for data analysis.

N-terminal peptides were identified by the presence of an N-terminal label or other blocking entity. Samples were compared using a Welch’s t-test with a Benjamini-Hochberg correction using a *P*-value of <0.05 for significance performed in Proteome Discoverer 2.2. Peptides had to be identified in more than 2 of 3 technical replicates in at least one group of samples for further consideration. Peptide annotation was performed using TopFIND (TopFINDER tool, http://clipserve.clip.ubc.ca/topfind/topfinder) to distinguish between internal and natural peptides (88-90). Data visualization utilized GraphPad Prism 9, PowerPoint, and Excel (Microsoft, 2013) and peptide positions were mapped on proteins based on their structure using uniprot.org annotations.

### Interference reflection microcopy (IRM)

Parental RPE1 or D12 cells, with or without siRNA treatment were seeded onto glass-bottom dishes (MatTek, catalog no. P35G-0-14-C) in phenol-red free medium and imaged at 30 min or 24 hrs after seeding. Live imaging was carried out for 2 h at 1 min intervals in a climate-controlled environmental chamber, using a Leica SP8 confocal microscope (Leica Microsystems). Images were acquired using a 40X/1.25 oil immersion objective at zoom of 1.275, 488 nm excitation with a PMT detector collecting window of 460 nm-520 nm. Histograms from each cell were used to quantify focal adhesions with the lowest brightness levels (intensity <40) representative of strong focal adhesions (distance between the cell and the plate <2.5 Å).

### Imaging of collagen proteolysis

8-well chamber slides were coated with 25 μg/ml DQ collagen type I (Invitrogen, catalog no. D12060) containing a 1:11 ratio of DQ-conjugated to non-conjugated bovine collagen I for 2 h at room temperature and seeded with parental RPE-1 or D12 cells followed by culture for 24 hours in serum-free DMEM+F12 culture medium. A coated culture well without seeded cells was used as a control to determine basal (quenched) fluorescence signal of DQ collagen type I. 24 h after seeding, culture medium was aspirated and the cell layer was washed once with PBS and fixed with 4% paraformaldehyde (PFA) in phosphate buffered saline (PBS) containing 0.1% Tween-20 (PBST), for 10 minutes with agitation. The fixed cell layer was washed three times using PBST and mounted overnight using ProLong Gold mounting medium containing DAPI (Invitrogen, P36931). Unquenched fluorescein DQ-collagen signal was imaged using a Zeiss Axioplan microscope equipped with a Leica DM6200 camera.

### Imaging of cell-surface MT1-MMP and quantitative assessment of cilia

Parental RPE-1 and D12 cells were seeded in 8-well chamber glass slides for 24 h and were treated with control siRNA or *MMP14* siRNA using Lipofectamine RNAi MAX transfection reagent overnight. Culture medium was aspirated and the cell layer was washed with sterile PBS and the cells were cultured in serum free DMEM-F12 culture medium for induction of cilia. For MT1-MMP overexpression experiments, 8-chamber glass slides were coated with 5μg/ml vitronectin (R&D systems, 2308-VN) for 2 hours at room temperature prior to being seeded with parental RPE-1 cells. After 24 h culture, the cells were transfected with 100 ng/well of pcDNA 3.1 Myc/His empty vector or MT1-MMP plasmid DNA using PEI max transfection reagent (Polysciences, catalog no. 24765). After 6 hours of treatment, cells were washed with PBS and cultured in serum-free DMEM-F12 medium for 24 hours. In both knockdown and overexpression experiments, the cells were harvested after 24 h by washing once with PBS followed by fixation in 4% PFA in PBST, for 15 minutes. Primary cilia were visualized by staining with anti-acetylated alpha tubulin (1:400, Invitrogen, catalog no. 32-2700) and imaged using a Zeiss Axioplan microcope equipped with a Leica DMC6200 camera. Cilium length and percentage was quantified using NIH Image J Fiji software as previously described (12). MT1-MMP immunostaining was conducted on cells grown in 8-chamber slides and fixed with 4% PFA in PBS (without 0.l% Tween-20), utilizing a 1:400 dilution of the rabbit monoclonal MT1-MMP antibody (EP1264Y) against the catalytic domain (Abcam, catalog no. ab51074). Blocking, primary and secondary antibody staining and washing steps were conducted under detergent-free conditions. Prior to coverslipping and mounting with the ProLong Gold mounting medium containing DAPI (Invitrogen, P36931), the cells were washed with PBST for 15 minutes to facilitate permeability for nuclear staining with DAPI.

### Quantitative reverse transcriptase PCR

Real time quantitative PCR (RT-qPCR) was performed as described in (add Nandadasa et al., (Matrix Biol. 2021). In brief, 2 μg of total RNA harvested from parental RPE-1 and D12 cells utilizing the TRIzol reagent (ThermoFisher Scientific, 15596026) was used for cDNA synthesis (Applied Biosystems, Thermo Fisher Scientific4368814). *GAPDH* expression was used for normalizing relative gene expression calculated by the ΔΔCt quantification method using a CFX96 Touch Real-Time PCR detection system (Bio-Rad Laboratories). An unpaired, two-tailed Student t-test was used to calculate statistical significance. The following primer pairs were used for RT-qPCR analysis. *GAPDH*-F 5’-AGCCTCAAGATCATCAGCAATG-3’; *GAPDH*-R 5’-CTTCCACGATACCAAAGTTGTCAT-3’, *MMP14*-F 5’-CCTTGGACTGTCAGGAATGAGG-3’; *MMP14*-R 5’-TTCTCCGTGTCCATCCACTGGT-3’, *MMP2*-F 5’-AGCGAGTGGATGCCGCCTTTAA-3’; *MMP2*-R 5’-CATTCCAGGCATCTGCGATGAG-3’.

## Data availability

The mass spectrometry proteomics data have been deposited to the ProteomeXchange Consortium via the PRIDE partner repository (91) with the dataset identifier PXD036612 and 10.6019/PXD036612. Reviewer account details are: **Username:**; **Password:** u9qk5Bpm

## Abbreviations

ADAMTS: A disintegrin-like and metalloproteinase domain with thrombospondin type 1 repeats.
ECM: Extracellular matrix
IRM: Interference reflection microscopy
MMP: Matrix metalloproteinase
MT1-MMP: Membrane type 1-matrix metalloproteinase
TAILS: Terminal amine isotopic labeling of substrates

## Acknowledgements

This work was supported by the Allen Distinguished Investigator Program, through support made by The Paul G. Allen Frontiers Group and the American Heart Association (Award 17DIA33820024 to S.S.A.), NIH R01DK126804-01A1 (to S.N.) and the Worcester Foundation award (to S.N). The Fusion Lumos instrument was purchased via an NIH shared instrument grant, 1S10OD023436-01.

## Notes

### Competing Interest Statement

The authors have declared no competing interest.

## References

1. Lichtenthaler, S. F., Lemberg, M. K., and Fluhrer, R. (2018) Proteolytic ectodomain shedding of membrane proteins in mammals-hardware, concepts, and recent developments. Embo j 37

2. Mead, T. J., and Apte, S. S. (2018) ADAMTS proteins in human disorders. Matrix Biol 71-72, 225–239

3. Yu, W. H., and Woessner, J. F., Jr. (2000) Heparan sulfate proteoglycans as extracellular docking molecules for matrilysin (matrix metalloproteinase 7). J Biol Chem 275, 4183–4191

4. Kuno, K., and Matsushima, K. (1998) ADAMTS-1 protein anchors at the extracellular matrix through the thrombospondin type I motifs and its spacing region. J Biol Chem 273, 13912–13917

5. Cain, S. A., Mularczyk, E. J., Singh, M., Massam-Wu, T., and Kielty, C. M. (2016) ADAMTS-10 and -6 differentially regulate cell-cell junctions and focal adhesions. Sci Rep 6, 35956

6. Rodriguez-Manzaneque, J. C., Carpizo, D., Plaza-Calonge Mdel, C., Torres-Collado, A. X., Thai, S. N., Simons, M., Horowitz, A., and Iruela-Arispe, M. L. (2009) Cleavage of syndecan-4 by ADAMTS1 provokes defects in adhesion. Int J Biochem Cell Biol 41, 800–810

7. Clark, M. E., Kelner, G. S., Turbeville, L. A., Boyer, A., Arden, K. C., and Maki, R. A. (2000) ADAMTS9, a novel member of the ADAM-TS/ metallospondin gene family. Genomics 67, 343–350.

8. Somerville, R. P., Longpre, J. M., Jungers, K. A., Engle, J. M., Ross, M., Evanko, S., Wight, T. N., Leduc, R., and Apte, S. S. (2003) Characterization of ADAMTS-9 and ADAMTS-20 as a distinct ADAMTS subfamily related to Caenorhabditis elegans GON-1. J Biol Chem 278, 9503–9513

9. Jungers, K. A., Le Goff, C., Somerville, R. P., and Apte, S. S. (2005) Adamts9 is widely expressed during mouse embryo development. Gene Expr Patterns 5, 609–617

10. Koo, B. H., Coe, D. M., Dixon, L. J., Somerville, R. P., Nelson, C. M., Wang, L. W., Young, M. E., Lindner, D. J., and Apte, S. S. (2010) ADAMTS9 is a cell-autonomously acting, anti-angiogenic metalloprotease expressed by microvascular endothelial cells. Am J Pathol 176, 1494–1504

11. McCulloch, D. R., Nelson, C. M., Dixon, L. J., Silver, D. L., Wylie, J. D., Lindner, V., Sasaki, T., Cooley, M. A., Argraves, W. S., and Apte, S. S. (2009) ADAMTS metalloproteases generate active versican fragments that regulate interdigital web regression. Dev Cell 17, 687–698

12. Nandadasa, S., Kraft, C. M., Wang, L. W., O’Donnell, A., Patel, R., Gee, H. Y., Grobe, K., Cox, T. C., Hildebrandt, F., and Apte, S. S. (2019) Secreted metalloproteases ADAMTS9 and ADAMTS20 have a non-canonical role in ciliary vesicle growth during ciliogenesis. Nat Commun 10, 953

13. Nandadasa, S., Nelson, C. M., and Apte, S. S. (2015) ADAMTS9-Mediated Extracellular Matrix Dynamics Regulates Umbilical Cord Vascular Smooth Muscle Differentiation and Rotation. Cell Rep 11, 1519–1528

14. Mead, T. J., Du, Y., Nelson, C. M., Gueye, N. A., Drazba, J., Dancevic, C. M., Vankemmelbeke, M., Buttle, D. J., and Apte, S. S. (2018) ADAMTS9-Regulated Pericellular Matrix Dynamics Governs Focal Adhesion-Dependent Smooth Muscle Differentiation. Cell Rep 23, 485–498

15. Benz, B. A., Nandadasa, S., Takeuchi, M., Grady, R. C., Takeuchi, H., LoPilato, R. K., Kakuda, S., Somerville, R. P., Apte, S. S., Haltiwanger, R. S., and Holdener, B. C. (2016) Genetic and biochemical evidence that gastrulation defects in Pofut2 mutants result from defects in ADAMTS9 secretion. Dev Biol

16. Dubail, J., Aramaki-Hattori, N., Bader, H. L., Nelson, C. M., Katebi, N., Matuska, B., Olsen, B. R., and Apte, S. S. (2014) A new Adamts9 conditional mouse allele identifies its non-redundant role in interdigital web regression. Genesis 52, 702–712

17. Dubail, J., Vasudevan, D., Wang, L. W., Earp, S. E., Jenkins, M. W., Haltiwanger, R. S., and Apte, S. S. (2016) Impaired ADAMTS9 secretion: A potential mechanism for eye defects in Peters Plus Syndrome. Sci Rep 6, 33974

18. Enomoto, H., Nelson, C., Somerville, R.P.T., Mielke, K., Dixon, L., Powell, K., Apte, S.S. (2010) Cooperation of two ADAMTS metalloproteases in closure of the mouse palate identifies a requirement for versican proteolysis in regulating palatal mesenchyme proliferation. Development 137, 4029–4038

19. Kern, C. B., Wessels, A., McGarity, J., Dixon, L. J., Alston, E., Argraves, W. S., Geeting, D., Nelson, C. M., Menick, D. R., and Apte, S. S. (2010) Reduced versican cleavage due to Adamts9 haploinsufficiency is associated with cardiac and aortic anomalies. Matrix Biol 29, 304–316

20. Blelloch, R., Anna-Arriola, S. S., Gao, D., Li, Y., Hodgkin, J., and Kimble, J. (1999) The gon-1 gene is required for gonadal morphogenesis in Caenorhabditis elegans. Dev Biol 216, 382–393

21. Blelloch, R., and Kimble, J. (1999) Control of organ shape by a secreted metalloprotease in the nematode Caenorhabditis elegans. Nature 399, 586–590

22. Ismat, A., Cheshire, A. M., and Andrew, D. J. (2013) The secreted AdamTS-A metalloprotease is required for collective cell migration. Development 140, 1981–1993

23. Carter, N. J., Roach, Z. A., Byrnes, M. M., and Zhu, Y. (2019) Adamts9 is necessary for ovarian development in zebrafish. Gen Comp Endocrinol 277, 130–140

24. Carver, J. J., He, Y., and Zhu, Y. (2021) Delay in primordial germ cell migration in adamts9 knockout zebrafish. Sci Rep 11, 8545

25. Choi, Y. J., Halbritter, J., Braun, D. A., Schueler, M., Schapiro, D., Rim, J. H., Nandadasa, S., Choi, W. I., Widmeier, E., Shril, S., Körber, F., Sethi, S. K., Lifton, R. P., Beck, B. B., Apte, S. S., Gee, H. Y., and Hildebrandt, F. (2019) Mutations of ADAMTS9 Cause Nephronophthisis-Related Ciliopathy. Am J Hum Genet 104, 45–54

26. Matsushita, H. B., Hiraide, T., Hayakawa, K., Okano, S., Nakashima, M., Saitsu, H., and Kato, M. (2021) Compound heterozygous ADAMTS9 variants in Joubert syndrome-related disorders without renal manifestation. Brain Dev

27. Rao, C., Foernzler, D., Loftus, S. K., Liu, S., McPherson, J. D., Jungers, K. A., Apte, S. S., Pavan, W. J., and Beier, D. R. (2003) A defect in a novel ADAMTS family member is the cause of the belted white-spotting mutation. Development 130, 4665–4672

28. Silver, D. L., Hou, L., Somerville, R., Young, M.E., Apte, S.S., Pavan, W.J. (2008) The secreted metalloprotease ADAMTS20 is required for melanoblast survival. PLoS Genet 4, 1–15

29. Wolf, Z. T., Brand, H. A., Shaffer, J. R., Leslie, E. J., Arzi, B., Willet, C. E., Cox, T. C., McHenry, T., Narayan, N., Feingold, E., Wang, X., Sliskovic, S., Karmi, N., Safra, N., Sanchez, C., Deleyiannis, F. W., Murray, J. C., Wade, C. M., Marazita, M. L., and Bannasch, D. L. (2015) Genome-wide association studies in dogs and humans identify ADAMTS20 as a risk variant for cleft lip and palate. PLoS Genet 11, e1005059

30. Nandadasa, S., Burin des Roziers, C., Koch, C., Tran-Lundmark, K., Dours-Zimmermann, M. T., Zimmermann, D. R., Valleix, S., and Apte, S. S. (2021) A new mouse mutant with cleavage-resistant versican and isoform-specific versican mutants demonstrate that proteolysis at the Glu(441)-Ala(442) peptide bond in the V1 isoform is essential for interdigital web regression. Matrix Biol Plus 10, 100064

31. Tharmarajah, G., Eckhard, U., Jain, F., Marino, G., Prudova, A., Urtatiz, O., Fuchs, H., de Angelis, M. H., Overall, C. M., and Van Raamsdonk, C. D. (2018) Melanocyte development in the mouse tail epidermis requires the Adamts9 metalloproteinase. Pigment Cell Melanoma Res 31, 693–707

32. Kleifeld, O., Doucet, A., auf dem Keller, U., Prudova, A., Schilling, O., Kainthan, R. K., Starr, A. E., Foster, L. J., Kizhakkedathu, J. N., and Overall, C. M. (2010) Isotopic labeling of terminal amines in complex samples identifies protein N-termini and protease cleavage products. Nat Biotechnol 28, 281–288

33. Gifford, V., and Itoh, Y. (2019) MT1-MMP-dependent cell migration: proteolytic and non-proteolytic mechanisms. Biochem Soc Trans 47, 811–826

34. Nandadasa, S., O’Donnell, A., Murao, A., Yamaguchi, Y., Midura, R. J., Olson, L., and Apte, S. S. (2021) The versican-hyaluronan complex provides an essential extracellular matrix niche for Flk1(+) hematoendothelial progenitors. Matrix Biol

35. Yamagata, M., and Kimata, K. (1994) Repression of a malignant cell-substratum adhesion phenotype by inhibiting the production of the anti-adhesive proteoglycan, PG-M/versican. J Cell Sci 107, 2581–2590

36. Yamagata, M., Saga, S., Kato, M., Bernfield, M., and Kimata, K. (1993) Selective distributions of proteoglycans and their ligands in pericellular matrix of cultured fibroblasts. Implications for their roles in cell-substratum adhesion. J Cell Sci 106, 55–65

37. Yamagata, M., Suzuki, S., Akiyama, S. K., Yamada, K. M., and Kimata, K. (1989) Regulation of cell-substrate adhesion by proteoglycans immobilized on extracellular substrates. J Biol Chem 264, 8012–8018

38. Mosher, D. F., Saksela, O., Keski-Oja, J., and Vaheri, A. (1977) Distribution of a major surface-associated glycoprotein, fibronectin, in cultures of adherent cells. J Supramol Struct 6, 551–557

39. Woods, A., Couchman, J. R., Johansson, S., and Höök, M. (1986) Adhesion and cytoskeletal organisation of fibroblasts in response to fibronectin fragments. Embo j 5, 665–670

40. Barr, V. A., and Bunnell, S. C. (2009) Interference reflection microscopy. Curr Protoc Cell Biol Chapter 4, Unit 4.23

41. Drazba, J., Liljelund, P., Smith, C., Payne, R., and Lemmon, V. (1997) Growth cone interactions with purified cell and substrate adhesion molecules visualized by interference reflection microscopy. Brain Res Dev Brain Res 100, 183–197

42. Bause, E., and Hettkamp, H. (1979) Primary structural requirements for N-glycosylation of peptides in rat liver. FEBS Lett 108, 341–344

43. Remacle, A. G., Chekanov, A. V., Golubkov, V. S., Savinov, A. Y., Rozanov, D. V., and Strongin, A. Y. (2006) O-glycosylation regulates autolysis of cellular membrane type-1 matrix metalloproteinase (MT1-MMP). J Biol Chem 281, 16897–16905

44. Wu, Y. I., Munshi, H. G., Sen, R., Snipas, S. J., Salvesen, G. S., Fridman, R., and Stack, M. S. (2004) Glycosylation broadens the substrate profile of membrane type 1 matrix metalloproteinase. J Biol Chem 279, 8278–8289

45. Hotary, K., Allen, E., Punturieri, A., Yana, I., and Weiss, S. J. (2000) Regulation of cell invasion and morphogenesis in a three-dimensional type I collagen matrix by membrane-type matrix metalloproteinases 1, 2, and 3. J Cell Biol 149, 1309–1323

46. Sato, H., Takino, T., Okada, Y., Cao, J., Shinagawa, A., Yamamoto, E., and Seiki, M. (1994) A matrix metalloproteinase expressed on the surface of invasive tumour cells. Nature 370, 61–65

47. Holmbeck, K., Bianco, P., Caterina, J., Yamada, S., Kromer, M., Kuznetsov, S. A., Mankani, M., Robey, P. G., Poole, A. R., Pidoux, I., Ward, J. M., and Birkedal-Hansen, H. (1999) MT1-MMP-deficient mice develop dwarfism, osteopenia, arthritis, and connective tissue disease due to inadequate collagen turnover. Cell 99, 81–92

48. Lehti, K., Allen, E., Birkedal-Hansen, H., Holmbeck, K., Miyake, Y., Chun, T. H., and Weiss, S. J. (2005) An MT1-MMP-PDGF receptor-beta axis regulates mural cell investment of the microvasculature. Genes Dev 19, 979–991

49. Oblander, S. A., Zhou, Z., Gálvez, B. G., Starcher, B., Shannon, J. M., Durbeej, M., Arroyo, A. G., Tryggvason, K., and Apte, S. S. (2005) Distinctive functions of membrane type 1 matrix-metalloprotease (MT1-MMP or MMP-14) in lung and submandibular gland development are independent of its role in pro-MMP-2 activation. Dev Biol 277, 255–269

50. Zhou, Z., Apte, S. S., Soininen, R., Cao, R., Baaklini, G. Y., Rauser, R. W., Wang, J., Cao, Y., and Tryggvason, K. (2000) Impaired endochondral ossification and angiogenesis in mice deficient in membrane-type matrix metalloproteinase I. Proc Natl Acad Sci U S A 97, 4052–4057

51. Taylor, S. H., Yeung, C. Y., Kalson, N. S., Lu, Y., Zigrino, P., Starborg, T., Warwood, S., Holmes, D. F., Canty-Laird, E. G., Mauch, C., and Kadler, K. E. (2015) Matrix metalloproteinase 14 is required for fibrous tissue expansion. Elife 4, e09345

52. Wang, L. W., Nandadasa, S., Annis, D. S., Dubail, J., Mosher, D. F., Willard, B. B., and Apte, S. S. (2019) A disintegrin-like and metalloproteinase domain with thrombospondin type 1 motif 9 (ADAMTS9) regulates fibronectin fibrillogenesis and turnover. J Biol Chem 294, 9924–9936

53. Hernandez-Barrantes, S., Toth, M., Bernardo, M. M., Yurkova, M., Gervasi, D. C., Raz, Y., Sang, Q. A., and Fridman, R. (2000) Binding of active (57 kDa) membrane type 1-matrix metalloproteinase (MT1-MMP) to tissue inhibitor of metalloproteinase (TIMP)-2 regulates MT1-MMP processing and pro-MMP-2 activation. J Biol Chem 275, 12080–12089

54. Toth, M., Hernandez-Barrantes, S., Osenkowski, P., Bernardo, M. M., Gervasi, D. C., Shimura, Y., Meroueh, O., Kotra, L. P., Gálvez, B. G., Arroyo, A. G., Mobashery, S., and Fridman, R. (2002) Complex pattern of membrane type 1 matrix metalloproteinase shedding. Regulation by autocatalytic cells surface inactivation of active enzyme. J Biol Chem 277, 26340–26350

55. Cho, J. A., Osenkowski, P., Zhao, H., Kim, S., Toth, M., Cole, K., Aboukameel, A., Saliganan, A., Schuger, L., Bonfil, R. D., and Fridman, R. (2008) The inactive 44-kDa processed form of membrane type 1 matrix metalloproteinase (MT1-MMP) enhances proteolytic activity via regulation of endocytosis of active MT1-MMP. J Biol Chem 283, 17391–17405

56. Osenkowski, P., Toth, M., and Fridman, R. (2004) Processing, shedding, and endocytosis of membrane type 1-matrix metalloproteinase (MT1-MMP). J Cell Physiol 200, 2–10

57. Takino, T., Watanabe, Y., Matsui, M., Miyamori, H., Kudo, T., Seiki, M., and Sato, H. (2006) Membrane-type 1 matrix metalloproteinase modulates focal adhesion stability and cell migration. Exp Cell Res 312, 1381–1389

58. Woskowicz, A. M., Weaver, S. A., Shitomi, Y., Ito, N., and Itoh, Y. (2013) MT-LOOP-dependent localization of membrane type I matrix metalloproteinase (MT1-MMP) to the cell adhesion complexes promotes cancer cell invasion. J Biol Chem 288, 35126–35137

59. Wang, Y., and McNiven, M. A. (2012) Invasive matrix degradation at focal adhesions occurs via protease recruitment by a FAK-p130Cas complex. J Cell Biol 196, 375–385

60. Grafinger, O. R., Gorshtein, G., Stirling, T., Geddes-McAlister, J., and Coppolino, M. G. (2021) Inhibition of β1 integrin induces its association with MT1-MMP and decreases MT1-MMP internalization and cellular invasiveness. Cell Signal 83, 109984

61. Mori, H., Lo, A. T., Inman, J. L., Alcaraz, J., Ghajar, C. M., Mott, J. D., Nelson, C. M., Chen, C. S., Zhang, H., Bascom, J. L., Seiki, M., and Bissell, M. J. (2013) Transmembrane/cytoplasmic, rather than catalytic, domains of Mmp14 signal to MAPK activation and mammary branching morphogenesis via binding to integrin β1. Development 140, 343–352

62. Deryugina, E. I., Ratnikov, B. I., Postnova, T. I., Rozanov, D. V., and Strongin, A. Y. (2002) Processing of integrin alpha(v) subunit by membrane type 1 matrix metalloproteinase stimulates migration of breast carcinoma cells on vitronectin and enhances tyrosine phosphorylation of focal adhesion kinase. J Biol Chem 277, 9749–9756

63. Gálvez, B. G., Matías-Román, S., Yáñez-Mó, M., Sánchez-Madrid, F., and Arroyo, A. G. (2002) ECM regulates MT1-MMP localization with beta1 or alphavbeta3 integrins at distinct cell compartments modulating its internalization and activity on human endothelial cells. J Cell Biol 159, 509–521

64. Ratnikov, B. I., Rozanov, D. V., Postnova, T. I., Baciu, P. G., Zhang, H., DiScipio, R. G., Chestukhina, G. G., Smith, J. W., Deryugina, E. I., and Strongin, A. Y. (2002) An alternative processing of integrin alpha(v) subunit in tumor cells by membrane type-1 matrix metalloproteinase. J Biol Chem 277, 7377–7385

65. Gonzalo, P., Guadamillas, M. C., Hernandez-Riquer, M. V., Pollan, A., Grande-Garcia, A., Bartolome, R. A., Vasanji, A., Ambrogio, C., Chiarle, R., Teixido, J., Risteli, J., Apte, S. S., del Pozo, M. A., and Arroyo, A. G. (2010) MT1-MMP is required for myeloid cell fusion via regulation of Rac1 signaling. Dev Cell 18, 77–89

66. Gonzalo, P., Moreno, V., Gálvez, B. G., and Arroyo, A. G. (2010) MT1-MMP and integrins: Hand-to-hand in cell communication. Biofactors 36, 248–254

67. Martin, C. E., Murray, A. S., Sala-Hamrick, K. E., Mackinder, J. R., Harrison, E. C., Lundgren, J. G., Varela, F. A., and List, K. (2021) Posttranslational modifications of serine protease TMPRSS13 regulate zymogen activation, proteolytic activity, and cell surface localization. J Biol Chem 297, 101227

68. Vandooren, J., Pereira, R. V. S., Ugarte-Berzal, E., Rybakin, V., Noppen, S., Stas, M. R., Bernaerts, E., Ganseman, E., Metzemaekers, M., Schols, D., Proost, P., and Opdenakker, G. (2021) Internal Disulfide Bonding and Glycosylation of Interleukin-7 Protect Against Proteolytic Inactivation by Neutrophil Metalloproteinases and Serine Proteases. Front Immunol 12, 701739

69. Wang, S., Foster, S. R., Sanchez, J., Corcilius, L., Larance, M., Canals, M., Stone, M. J., and Payne, R. J. (2021) Glycosylation Regulates N-Terminal Proteolysis and Activity of the Chemokine CCL14. ACS Chem Biol 16, 973–981

70. Madzharova, E., Kastl, P., Sabino, F., and Auf dem Keller, U. (2019) Post-Translational Modification-Dependent Activity of Matrix Metalloproteinases. Int J Mol Sci 20

71. Guimier, A., Gabriel, G. C., Bajolle, F., Tsang, M., Liu, H., Noll, A., Schwartz, M., El Malti, R., Smith, L. D., Klena, N. T., Jimenez, G., Miller, N. A., Oufadem, M., Moreau de Bellaing, A., Yagi, H., Saunders, C. J., Baker, C. N., Di Filippo, S., Peterson, K. A., Thiffault, I., Bole-Feysot, C., Cooley, L. D., Farrow, E. G., Masson, C., Schoen, P., Deleuze, J. F., Nitschké, P., Lyonnet, S., de Pontual, L., Murray, S. A., Bonnet, D., Kingsmore, S. F., Amiel, J., Bouvagnet, P., Lo, C. W., and Gordon, C. T. (2015) MMP21 is mutated in human heterotaxy and is required for normal left-right asymmetry in vertebrates. Nat Genet 47, 1260–1263

72. Perles, Z., Moon, S., Ta-Shma, A., Yaacov, B., Francescatto, L., Edvardson, S., Rein, A. J., Elpeleg, O., and Katsanis, N. (2015) A human laterality disorder caused by a homozygous deleterious mutation in MMP21. J Med Genet 52, 840–847

73. Yuan, Z. Z., Fan, L. L., Jiang, Z. C., Yang, Y. F., and Tan, Z. P. (2020) A Novel Nonsense MMP21 Variant Causes Dextrocardia and Congenital Heart Disease in a Han Chinese Patient. Front Cardiovasc Med 7, 582350

74. Jiang, Z., Zhou, J., Qin, X., Zheng, H., Gao, B., Liu, X., Jin, G., and Zhou, Z. (2020) MT1-MMP deficiency leads to defective ependymal cell maturation, impaired ciliogenesis, and hydrocephalus. JCI Insight 5

75. Holdener, B. C., Percival, C. J., Grady, R. C., Cameron, D. C., Berardinelli, S. J., Zhang, A., Neupane, S., Takeuchi, M., Jimenez-Vega, J. C., Uddin, S. M. Z., Komatsu, D. E., Honkanen, R., Dubail, J., Apte, S. S., Sato, T., Narimatsu, H., McClain, S. A., and Haltiwanger, R. S. (2019) ADAMTS9 and ADAMTS20 are differentially affected by loss of B3GLCT in mouse model of Peters plus syndrome. Hum Mol Genet 28, 4053–4066

76. auf dem Keller, U., Doucet, A., and Overall, C. M. (2007) Protease research in the era of systems biology. Biol Chem 388, 1159–1162

77. Chan, K. M., Wong, H. L., Jin, G., Liu, B., Cao, R., Cao, Y., Lehti, K., Tryggvason, K., and Zhou, Z. (2012) MT1-MMP inactivates ADAM9 to regulate FGFR2 signaling and calvarial osteogenesis. Dev Cell 22, 1176–1190

78. Albrechtsen, R., Kveiborg, M., Stautz, D., Vikeså, J., Noer, J. B., Kotzsh, A., Nielsen, F. C., Wewer, U. M., and Fröhlich, C. (2013) ADAM12 redistributes and activates MMP-14, resulting in gelatin degradation, reduced apoptosis and increased tumor growth. J Cell Sci 126, 4707–4720

79. Zhou, Z., Apte, S. S., Soininen, R., Cao, R., Baaklini, G. Y., Rauser, R. W., Wang, J., Cao, Y., and Tryggvason, K. (2000) Impaired endochondral ossification and angiogenesis in mice deficient in membrane-type matrix metalloproteinase I. Proc Natl Acad Sci U S A 97, 4052–4057.

80. Fritsche, L. G., Chen, W., Schu, M., Yaspan, B. L., Yu, Y., Thorleifsson, G., Zack, D. J., Arakawa, S., Cipriani, V., Ripke, S., Igo, R. P., Jr., Buitendijk, G. H., Sim, X., Weeks, D. E., Guymer, R. H., Merriam, J. E., Francis, P. J., Hannum, G., Agarwal, A., Armbrecht, A. M., Audo, I., Aung, T., Barile, G. R., Benchaboune, M., Bird, A. C., Bishop, P. N., Branham, K. E., Brooks, M., Brucker, A. J., Cade, W. H., Cain, M. S., Campochiaro, P. A., Chan, C. C., Cheng, C. Y., Chew, E. Y., Chin, K. A., Chowers, I., Clayton, D. G., Cojocaru, R., Conley, Y. P., Cornes, B. K., Daly, M. J., Dhillon, B., Edwards, A. O., Evangelou, E., Fagerness, J., Ferreyra, H. A., Friedman, J. S., Geirsdottir, A., George, R. J., Gieger, C., Gupta, N., Hagstrom, S. A., Harding, S. P., Haritoglou, C., Heckenlively, J. R., Holz, F. G., Hughes, G., Ioannidis, J. P., Ishibashi, T., Joseph, P., Jun, G., Kamatani, Y., Katsanis, N. C N. K., Khan, J. C., Kim, I. K., Kiyohara, Y., Klein, B. E., Klein, R., Kovach, J. L., Kozak, I., Lee, C. J., Lee, K. E., Lichtner, P., Lotery, A. J., Meitinger, T., Mitchell, P., Mohand-Said, S., Moore, A. T., Morgan, D. J., Morrison, M. A., Myers, C. E., Naj, A. C., Nakamura, Y., Okada, Y., Orlin, A., Ortube, M. C., Othman, M. I., Pappas, C., Park, K. H., Pauer, G. J., Peachey, N. S., Poch, O., Priya, R. R., Reynolds, R., Richardson, A. J., Ripp, R., Rudolph, G., Ryu, E., Sahel, J. A., Schaumberg, D. A., Scholl, H. P., Schwartz, S. G., Scott, W. K., Shahid, H., Sigurdsson, H., Silvestri, G., Sivakumaran, T. A., Smith, R. T., Sobrin, L., Souied, E. H., Stambolian, D. E., Stefansson, H., Sturgill-Short, G. M., Takahashi, A., Tosakulwong, N., Truitt, B. J., Tsironi, E. E., Uitterlinden, A. G., van Duijn, C. M., Vijaya, L., Vingerling, J. R., Vithana, E. N., Webster, A. R., Wichmann, H. E., Winkler, T. W., Wong, T. Y., Wright, A. F., Zelenika, D., Zhang, M., Zhao, L., Zhang, K., Klein, M. L., Hageman, G. S., Lathrop, G. M., Stefansson, K., Allikmets, R., Baird, P. N., Gorin, M. B., Wang, J. J., Klaver, C. C., Seddon, J. M., Pericak-Vance, M. A., Iyengar, S. K., Yates, J. R., Swaroop, A., Weber, B. H., Kubo, M., Deangelis, M. M., Leveillard, T., Thorsteinsdottir, U., Haines, J. L., Farrer, L. A., Heid, I. M., Abecasis, G. R., and Consortium, A. M. D. G. (2013) Seven new loci associated with age-related macular degeneration. Nat Genet 45, 433-439, 439e431-432

81. Helisalmi, S., Immonen, I., Losonczy, G., Resch, M. D., Benedek, S., Balogh, I., Papp, A., Berta, A., Uusitupa, M., Hiltunen, M., and Kaarniranta, K. (2014) ADAMTS9 locus associates with increased risk of wet AMD. Acta Ophthalmol 92, e410

82. Whitmore, S. S., Braun, T. A., Skeie, J. M., Haas, C. M., Sohn, E. H., Stone, E. M., Scheetz, T. E., and Mullins, R. F. (2013) Altered gene expression in dry age-related macular degeneration suggests early loss of choroidal endothelial cells. Mol Vis 19, 2274–2297

83. van Zyl, T., Yan, W., McAdams, A., Peng, Y. R., Shekhar, K., Regev, A., Juric, D., and Sanes, J. R. (2020) Cell atlas of aqueous humor outflow pathways in eyes of humans and four model species provides insight into glaucoma pathogenesis. Proc Natl Acad Sci U S A 117, 10339–10349

84. Leightner, A. C., Hommerding, C. J., Peng, Y., Salisbury, J. L., Gainullin, V. G., Czarnecki, P. G., Sussman, C. R., and Harris, P. C. (2013) The Meckel syndrome protein meckelin (TMEM67) is a key regulator of cilia function but is not required for tissue planar polarity. Hum Mol Genet 22, 2024–2040

85. Baala, L., Romano, S., Khaddour, R., Saunier, S., Smith, U. M., Audollent, S., Ozilou, C., Faivre, L., Laurent, N., Foliguet, B., Munnich, A., Lyonnet, S., Salomon, R., Encha-Razavi, F., Gubler, M. C., Boddaert, N., de Lonlay, P., Johnson, C. A., Vekemans, M., Antignac, C., and Attie-Bitach, T. (2007) The Meckel-Gruber syndrome gene, MKS3, is mutated in Joubert syndrome. Am J Hum Genet 80, 186–194

86. Smith, U. M., Consugar, M., Tee, L. J., McKee, B. M., Maina, E. N., Whelan, S., Morgan, N. V., Goranson, E., Gissen, P., Lilliquist, S., Aligianis, I. A., Ward, C. J., Pasha, S., Punyashthiti, R., Malik Sharif, S., Batman, P. A., Bennett, C. P., Woods, C. G., McKeown, C., Bucourt, M., Miller, C. A., Cox, P., Algazali, L., Trembath, R. C., Torres, V. E., Attie-Bitach, T., Kelly, D. A., Maher, E. R., Gattone, V. H., 2nd, Harris, P. C., and Johnson, C. A. (2006) The transmembrane protein meckelin (MKS3) is mutated in Meckel-Gruber syndrome and the wpk rat. Nat Genet 38, 191–196

87. Weaver, S. A., Wolters, B., Ito, N., Woskowicz, A. M., Kaneko, K., Shitomi, Y., Seiki, M., and Itoh, Y. (2014) Basal localization of MT1-MMP is essential for epithelial cell morphogenesis in 3D collagen matrix. J Cell Sci 127, 1203–1213

88. Fortelny, N., Yang, S., Pavlidis, P., Lange, P. F., and Overall, C. M. (2015) Proteome TopFIND 3.0 with TopFINDer and PathFINDer: database and analysis tools for the association of protein termini to pre- and post-translational events. Nucleic Acids Res 43, D290–297

89. Lange, P. F., Huesgen, P. F., and Overall, C. M. (2012) TopFIND 2.0--linking protein termini with proteolytic processing and modifications altering protein function. Nucleic Acids Res 40, D351–361

90. Lange, P. F., and Overall, C. M. (2011) TopFIND, a knowledgebase linking protein termini with function. Nat Methods 8, 703–704

91. Perez-Riverol, Y., Bai, J., Bandla, C., García-Seisdedos, D., Hewapathirana, S., Kamatchinathan, S., Kundu, D. J., Prakash, A., Frericks-Zipper, A., Eisenacher, M., Walzer, M., Wang, S., Brazma, A., and Vizcaíno, J. A. (2022) The PRIDE database resources in 2022: a hub for mass spectrometry-based proteomics evidences. Nucleic Acids Res 50, D543–d552

