## Supplemental text and figures for "Degradomic identification of membrane type 1-matrix metalloproteinase (MT1-MMP/MMP14) as an ADAMTS9 and ADAMTS20 substrate"

**Content**:

The supplement contains one figure (Figure S1) and four movies (movies S1-S4)

**Figure S1**. D12 cells have significantly higher MT1-MMP proteolysis at a distinct site in the hinge.

(A) Cartoon of furin-processed MT1-MMP showing the cleavage site.

(B) MS2 spectrum of the peptide with an N-terminal iTRAQ label, whose sequence suggests the cleavage site.

(C) Quantitation of the peptides in B showing higher levels in D12 cells than in parental RPE-1 cells.

**Supplemental Movies**:

Movies S1-S2: Time-lapse videos taken over a 2 hour duration showing combined IRM and DIC microscopy of wild-type RPE-1 cells (Movie S1) and ADAMTS9-mutant D12 cells (Movie S2) taken 30 minutes after cell seeding. Cell adhesions on IRM are seen as dark areas superimposed on the cell boundaries visualized by DIC.

Movies S3-S4: The time-lapse videos were taken over a 2 h duration and show combined IRM and DIC microscopy of ADAMTS9-mutant D12 cells taken 24 hours after cell seeding. Cell adhesions on IRM are seen as dark areas superimposed on the cell boundaries visualized by DIC. Movie S3 and S4 show cells transfected with a control siRNA or MMP14 siRNA respectively

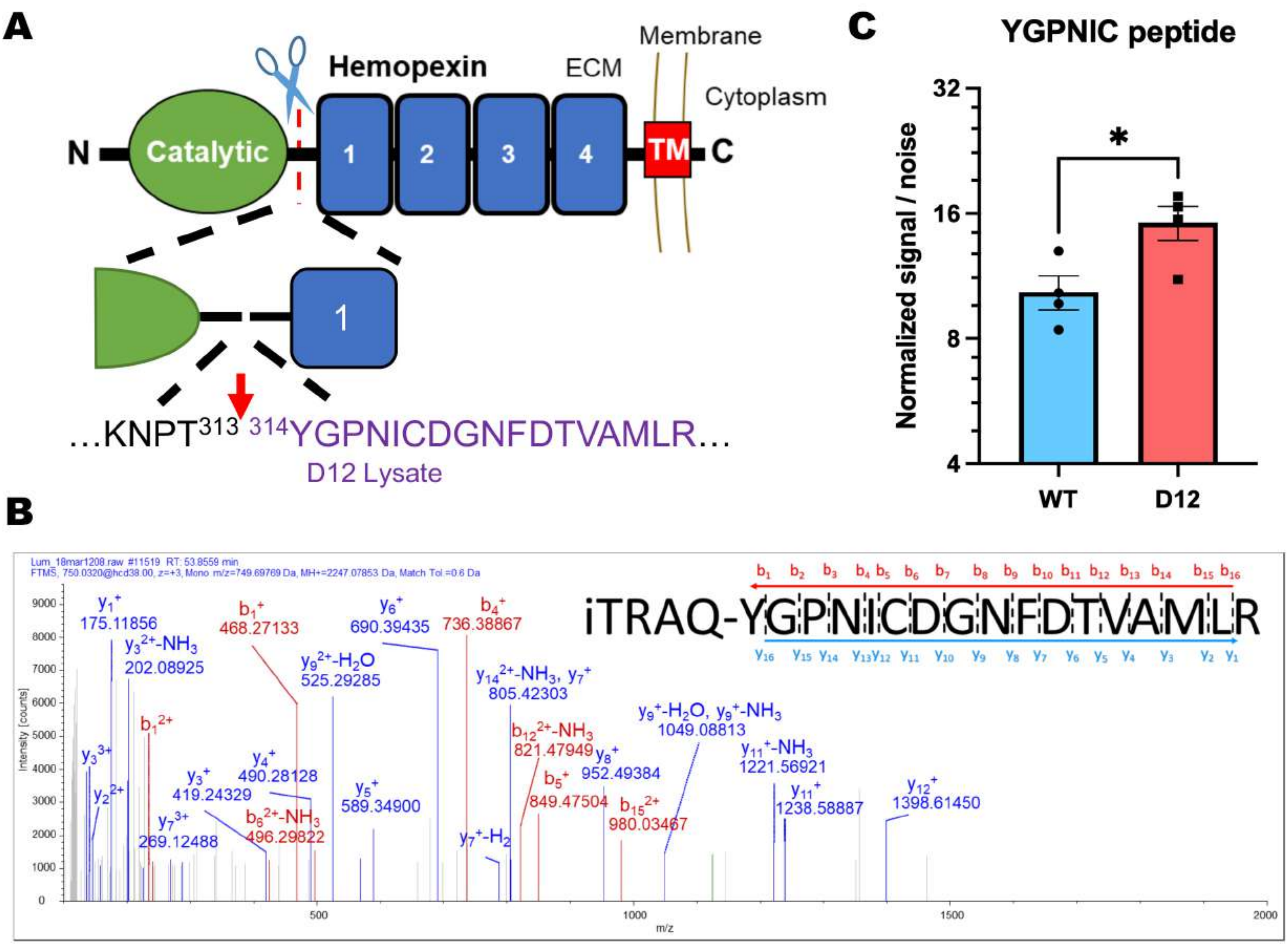
A
Hemopexin
ECM
Membrane
Cytoplasm
N
Catalytic
1
2
3
4
TM
C
1
…KNPT313 314YGPNICDGNFDTVAMLR…
D12 Lysate
B
Lum_18mar1208.raw #11519 RT: 53.8559 min
FTMS, 750.0320@hcd38.00, z=+3, Mono m/z=749.69769 Da, MH+=2247.07853 Da, Match Tol.=0.6 Da
iTRAQ-YGPNICDGNFDTVAMLR
y1+ 175.11856
y32+-NH3 202.08925
b1+ 468.27133
y92+-H2O 525.29285
y6+ 690.39435
b4+ 736.38867
y142+-NH3, y7+ 805.42303
b1 2+
b122+-NH3 821.47949
y9+-H2O, y9+-NH3 1049.08813
y33+
y4+ 490.28128
y3+ 419.24329
y8+ 952.49384
y11+-NH3 1221.56921
y22+
y5+ 589.34900
b5+ 849.47504
y11+ 1238.58887
y12+ 1398.61450
y73+ 269.12488
b62+-NH3 496.29822
b152+ 980.03467
y7+-H2
Intensity [counts]
m/z
C
YGPNIC peptide
Normalized signal / noise
32
16
8
4
*
WT
D12

Figure-S1
